# An Interpretable Model of pre-mRNA Splicing for Animal and Plant Genes

**DOI:** 10.1101/2023.12.29.573658

**Authors:** Kayla McCue, Christopher B. Burge

## Abstract

Pre-mRNA splicing is a fundamental step in gene expression, conserved across eukaryotes, in which the spliceosome recognizes motifs at the 3’ and 5’ splice sites (SS), excises introns and ligates exons. SS recognition and pairing is often influenced by splicing regulatory factors (SRFs) that bind to splicing regulatory elements (SREs). Several families of sequence-specific SRFs are known to be similarly ancient. Here, we describe SMsplice, a fully interpretable model of pre-mRNA splicing that combines new models of core SS motifs, SREs, and exonic and intronic length preferences. We learn models the predict SS locations with 83-86% accuracy in fish, insects and plants, and about 70% in mammals. Learned SRE motifs include both known SRF binding motifs as well as novel motifs, and both classes are supported by genetic analyses. Our comparisons across species highlight similarities between non-mammals and a greater reliance on SREs in mammalian splicing, and increased reliance on intronic SREs in plant splicing.

## Introduction

The removal of intronic sequences from pre-mRNA transcripts, known as splicing, constitutes a key step in the maturation process of that transcript. Catalyzed by the spliceosome, splicing is widespread in eukaryotic organisms and essential for expression of many genes (Keren et al., 2010). The 5’ and 3’ splice site (SS) motifs and the branch point sequences (BPS) form the core sequence elements required for splicing. These motifs are recognized by components of the spliceosome in a process that pairs the 5’ and 3’SS to define the intron between them (Wahl et al., 2009). However, these motifs do not, by themselves, contain sufficient information to fully inform the splicing patterns observed in many organisms (Lim & Burge, 2001). Instead, splicing is additionally affected by diverse splicing regulatory elements (SREs), which are recognized by a wide array of protein splicing factors (SFs), many deeply conserved in evolution (Busch & Hertel, 2012).

Large-scale cell-based screening has been used to identify sequences that impact exon inclusion, intron inclusion or splice site usage (Rosenberg et al., 2015; Y. Wang et al., 2012, 2013; Z. Wang et al., 2004), with most of these studies defining specific sets of SREs. Other studies have used computational approaches followed by minigene validation experiments to identify exonic SREs (Fairbrother et al., 2002; Goren et al., 2006), and a great deal of mutational analysis has identified SREs active in specific exons or introns (Baeza-Centurion et al., 2019, 2020; Ke et al., 2018). Likewise, dozens of SFs have been extensively studied biochemically and genetically (Fu & Ares, 2014). In vitro binding preferences of RBPs, including many splicing factors, have been mapped (Dominguez et al., 2018; Ray et al., 2013), and tools for modeling binding have been developed (Jens et al., 2022; Rube et al., 2022). Recently, in vivo binding and splicing activity following RNAi knockdown have been assessed for dozens of splicing factors (Van Nostrand et al., 2020). SS motifs have also been explored experimentally as well, including a recent large-scale screen that measured the activity of all possible 5’SS sequences in three different exonic contexts (Wong et al., 2018). Various computational models focused on determining SS strength and classifying short sequences as 5’ or 3’ SS have also been developed, and were among the earliest models related to splicing (Cheng et al., 2019; Shapiro & Senapathy, 1987; Yeo & Burge, 2004).

Over the past two decades, more comprehensive models of splicing have been developed, predominantly focused on mammalian systems. An early approach showed that adding known exonic splicing enhancer (ESE) and especially exonic splicing silencer (ESS) elements to SS motifs improved the accuracy of SS prediction (Z. Wang et al., 2004). Other models of splicing have been focused more on the prediction of splice-altering mutations, or the percent spliced in (PSI) for cassette (alternatively spliced) exons. These models use a variety of features to make their predictions, for instance, HAL predicts the change in PSI for a cassette exon following mutation based on the hexameric compositions of the wild type and variant sequences (Rosenberg et al., 2015). There have also been efforts to define a comprehensive ‘splicing code’ of relevant *cis*-acting elements, with over one thousand features, including exon lengths and binding motifs for known SFs (Barash et al., 2010; Xiong et al., 2011, 2015). These features have then been used in Bayesian neural network models in order to predict splicing disruption caused by genetic variants and relative PSI values for cassette alternative exons in different tissues. Models have also been developed that more explicitly seek to determine the likelihood of a variant disrupting splicing (Jian et al., 2014; Mort et al., 2014).

More recently, extremely high predictive performance for splicing has been achieved via the application of deep neural network methods such as SpliceAI (Jaganathan et al., 2019), which predicts splice sites from large segments of flanking human genomic sequence (up to 10 kb). These methods produce “black box” models, which learn parameters that are essentially uninterpretable. In silico mutagenesis experiments can be used to interrogate which sequence regions are important for specific predictions (Jaganathan et al., 2019). A mutational scan of hundreds of regions identified the expected reliance on core SS motifs but failed to detect enrichment for known SRE motifs, suggesting that the genomic features learned by the model overlap only partially with those used in SS recognition by the spliceosome (Gupta et al., 2023).

“White box” models, in contrast to black box models, are designed to be readily interpretable. They allow users to understand how the inputs and parameters are used to reach the conclusions of the model. Therefore, we developed a white box model where the structure and parameters are directly inspired by different elements of the splicing process. The resulting interpretability allowed us to assess the relative importance of “structural” features such as exon and intron lengths, and to derive scores for short sequences as splicing regulatory elements (SREs) that we show capture many known regulatory motifs and identify other motifs which are predictive of function.

To this end, we assumed that recognition by the spliceosome primarily involves three key aspects: motifs of variable strength at the 5’ and 3’SS; SREs located near the SS; and SS pairing, including distance constraints and preferences. New models of SS strength incorporating triplet preferences in a maximum entropy framework were found to improve on previous non-neural models (Yeo & Burge, 2004). Here, SREs were assumed to act locally (e.g., within 80 or 100 nt of the SS) and additively to enhance or suppress usage of nearby SS. Combining these first two features yielded a simple model we call CASS, which may have applications as a measure of SS strength in local context. Finally, we assumed that pairing of SS by the spliceosome involves not only steric considerations that enforce a minimum intron length, but also preferences for specific exon and intron lengths that reflect processes of exon and intron definition (De Conti et al., 2013). These three features were modeled together in our SMsplice model, whose structure is closely related to a Hidden Semi-Markov model (HSMM), enabling use of a classic HSMM algorithm to identify the most likely splicing pattern in a transcript. To maintain tractability, other features known to play a role in the splicing of some or all introns were omitted, including the BPS motif, recursive splicing, RNA secondary structure of the pre-mRNA, and impacts of SREs at longer distances (Sibley et al., 2016). Despite these omissions, our model yielded moderately to highly accurate predictions in a variety of animals and a model plant, and yielded putative SRE motifs in each organism as well as insights into the relative importance of different features across evolution.

## Results

### Context-aware SS scores more than double the performance of MaxEnt scores alone

We sought to develop scores representing the potential of individual sequence positions to function as 5’ or 3’SS, considering first core SS motifs, and then the impact of nearby SREs (Figure 1A). For SS motif scores, we developed an updated version of the MaxEntScan method (Yeo & Burge, 2004), taking advantage of much larger training sets of high-quality splice sites and improved computational resources to develop more complex and accurate models. Our new version is a “third-order” model as it captures dependencies between triplets of positions rather than just the pairs of positions in the core motifs considered previously (Methods). We defined core motif regions as in the original model, consisting of 9 nt at the 5’SS (–3 to +6), and 23 nt at the 3’SS (–20 to +3, including the polypyrimidine tract). The models return a log-odds score of a given sequence of length 9 or 23, based on the ratio of the probability of the sequence as a 5’SS or 3’SS divided by the probability of the sequence under a background model. This new version of the method yielded improved discrimination over the original, with a test set area under the receiver-operator characteristic curve (AUC) of 0.9982 compared to the original’s 0.9971 for 5’SS, and 0.9965 versus the original’s 0.9960 for the 3’SS model (Figure 1B). To use these models as SS predictors, we simply set a cutoff on the SS scores, and predicted as SS any position with score exceeding a threshold that maximized the F_1_ score, defined as the harmonic mean of precision and recall, on a training set of human genes (Methods). Application to a short human gene, *ZNF575,* is shown (Figure 1C).

**Figure 1:**
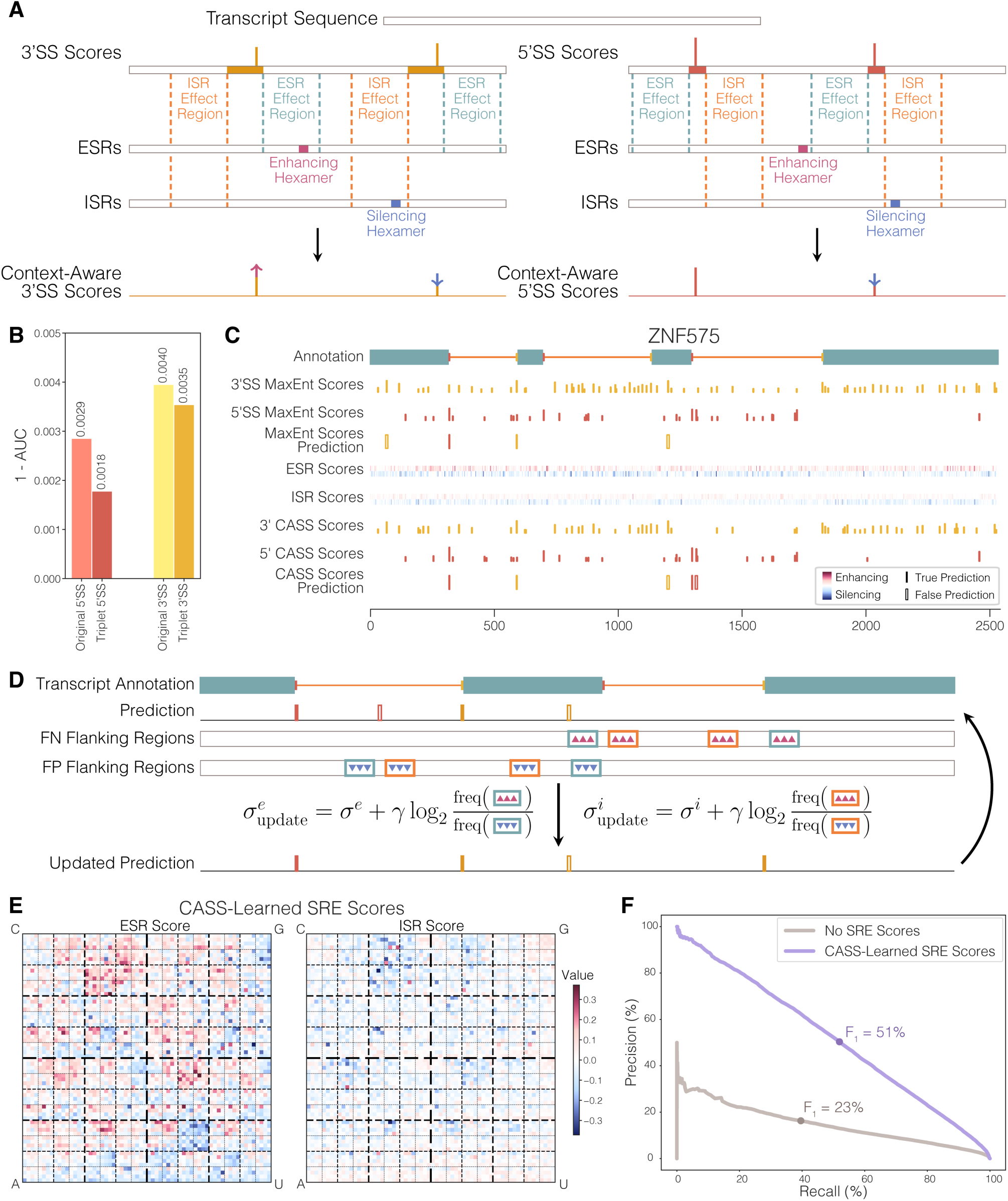
Context-Aware Splice Site Scores Improve Predictions. A) The CASS scores for an RNA sequence are determined by scoring every base in that sequence as a 5’SS and a 3’SS separately, treating the surrounding region as a potential SS motif, while independently scoring each hexamer in the sequence as an ESR and an ISR. Then, for each base its 3’ CASS score is the sum of its original 3’SS score, the ISR scores for the hexamers in the upstream ISR effect region, and the ESR scores for the hexamers in the downstream ESR effect region. The 5’ CASS score is similarly calculated, but with upstream ESR scores and downstream ISR scores. B) Smaller 1-AUC values for the updated MaxEnt methods compared with the originals show their improvement on classifying SS versus non-SS sequences. C) Illustration of the MaxEnt scores, SRE scores, CASS scores, and predictions on the gene *ZNF575* along with its annotation. For the 5’ and 3’SS, scores greater than zero are represented with height indicating the strength. For the ESR and ISR scores, positively scoring hexamers are in the top portion of the row and negatively scoring ones are in the bottom portion of the row, with intensity of color reflecting strength. For different predictions, solid lines represent correct predictions, hollow lines represent incorrect ones. D) Iterative learning of scores is performed by taking the frequency of hexamer appearances in regions flanking false positive and false negative predictions. Hexamers that appear frequently in exonic and intronic false positive regions compared to the respective false negative regions have their respective SRE scores decreased. Hexamers that appear frequently in exonic and intronic false negative regions compared to the respective false positive regions have their respective SRE scores increased. E) Chaos plots for CASS-learned SRE scores. For each hexamer, the first letter determines the overall quadrant in the heatmap, indicated at the corners. Then, the second letter of the hexamer, determines the subquadrant, and so on. The colors indicate the score for each hexamer. F) Precision-recall curves on the test set for predictions made using no SRE scores and the CASS-learned SRE scores. The dots represent the location on the curve determined from the cutoff learned on the training data, and the associated F_1_ score on the test data are indicated.

The activities of splice site motifs in human genes are known to be quite sensitive to the local sequence context, often being influenced by nearby SREs (Keren et al., 2010), which can act from exonic or intronic locations. Exonic SREs are typically designated exonic splicing enhancers (ESEs) when they promote use of upstream 3’SS/downstream 5’SS and/or inclusion of the exon in which they reside, and as exonic splicing silencers (ESSs) when they have the opposite effect on splicing. Similarly, intronic SREs that promote the use of downstream 3’SS or upstream 5’SS are known as intronic splicing enhancers (ISEs), while those that have the opposite effect are intronic splicing silencers (ISSs). To represent the behavior of SREs enhancing or silencing splicing in local regions, we defined an “ESR score” for each possible hexanucleotide (hexamer), representing its impact on the splicing of upstream 3’SS and downstream 5’SS, as well as an “ISR score” representing its impact on downstream 3’SS and upstream 5’SS. In essence, we treat hexamers with positive or negative ESR scores as akin to ESEs and ESSs, respectively, and those with positive or negative ISR scores as ISEs or ISSs. These scores, determined by a learning procedure describe below, are added to the “core SS scores” (from the MaxEnt procedure described above) of adjacent SS motifs as shown (Figure 1A). We refer to these SRE-modulated SS scores as context-aware SS (CASS) scores.

Mathematically, the CASS scores are defined by the formulas:

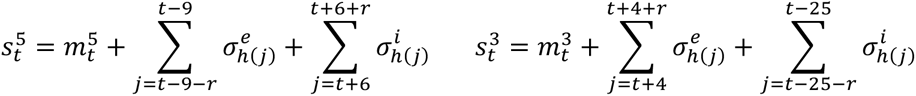

Here, *t* refers to the position of the base in question within the wider sequence and the superscripts 3 and 5 differentiate between the 3’ and 5’SS. So, 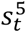 refers to the 5’ CASS score at position *t* of a sequence and 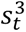 refers to the 3’ CASS score at that same position. The first terms in these equations, the *m* , are the core SS scores, and they have the same notation as the context-aware scores. For the summation terms, *r* indicates the range of sequence context considered and the bounds of the summation are chosen to avoid overlapping the sequence used to determine the SS scores. The value selected for *r* was 80, which proved optimal or near-optimal in a variety of tests in all organisms studied. The summands are the relevant SRE scores, the superscripts *e* and *i* differentiate between exonic and intronic context, and the subscript function *h*(*j*) is a hash function, indicating the index used for *σ*^’^ or *σ*, to get the value associated with the hexamer beginning at position *j* in the sequence.

ESR and ISR scores were learned as follows. Starting with the predictions made using our core SS motif models, we considered all positions that disagreed with the canonical annotation for the associated gene. We reasoned that these false positives (FP) and false negatives (FN) represented sequences where SREs were likely involved. An FP represents a strong SS motif which might require silencing elements in the flanking sequence, whereas an FN might require splicing enhancers nearby to promote its use. In particular, we took the flanking regions that would be incorporated into the CASS scores, splitting apart the exonic and intronic sides, and updated the SRE scores, encouraging the hexamers frequently present near FPs to be silencing and those frequently near FNs to be enhancing (Figure 1D, Methods). We then used these updated SRE scores to make new predictions on the training set and repeated this process until performance on the validation set began to decrease (Methods).

We refer to SRE scores calculated in this manner as the CASS-learned SRE scores. As the ESR and ISR scores are separate in the CASS model, we observed that, generally speaking, the ESR scores had greater magnitude, both positive and negative, than ISR scores (Figure 1E). Furthermore, we were able to quantify the benefit provided by the context in CASS scores. We applied the CASS framework to a test set of genes and, using the prediction cutoffs learned from the training set, calculated the F_1_ scores (Figure 1F). The ensuing change in performance was substantial, increasing from 23% with SS scores alone to 51% with CASS-learned SREs. Thus, more than half of the performance of the CASS model in human is attributable to the local SRE context of splice sites.

### SMsplice captures structural and sequence features of splicing and improves over CASS

CASS scores represent SS strength in local context but do not account for the pairing of splice sites across introns and/or exons that occurs in splicing. For instance, the CASS scores could predict a 3’SS upstream of the first plausible 5’SS in the gene (e.g., Figure 1C), though such a site could not participate in *cis*-splicing. Furthermore, there are known minimum intron lengths in human and other organisms below which splicing is not observed (Wieringa et al., 1984), and both exon and intron lengths can impact recognition by the spliceosome, often via exon or intron definition (De Conti et al., 2013). We therefore sought to make a more general model that could incorporate both splice site pairing and exon/intron length preferences, as well as the preferred numbers of introns per transcript. To this end, we developed a directed graphic structure which could describe the splicing pattern of an arbitrary transcript in terms of exons, introns, and splice sites beginning from the start of the transcript (Figure 2A).

**Figure 2:**
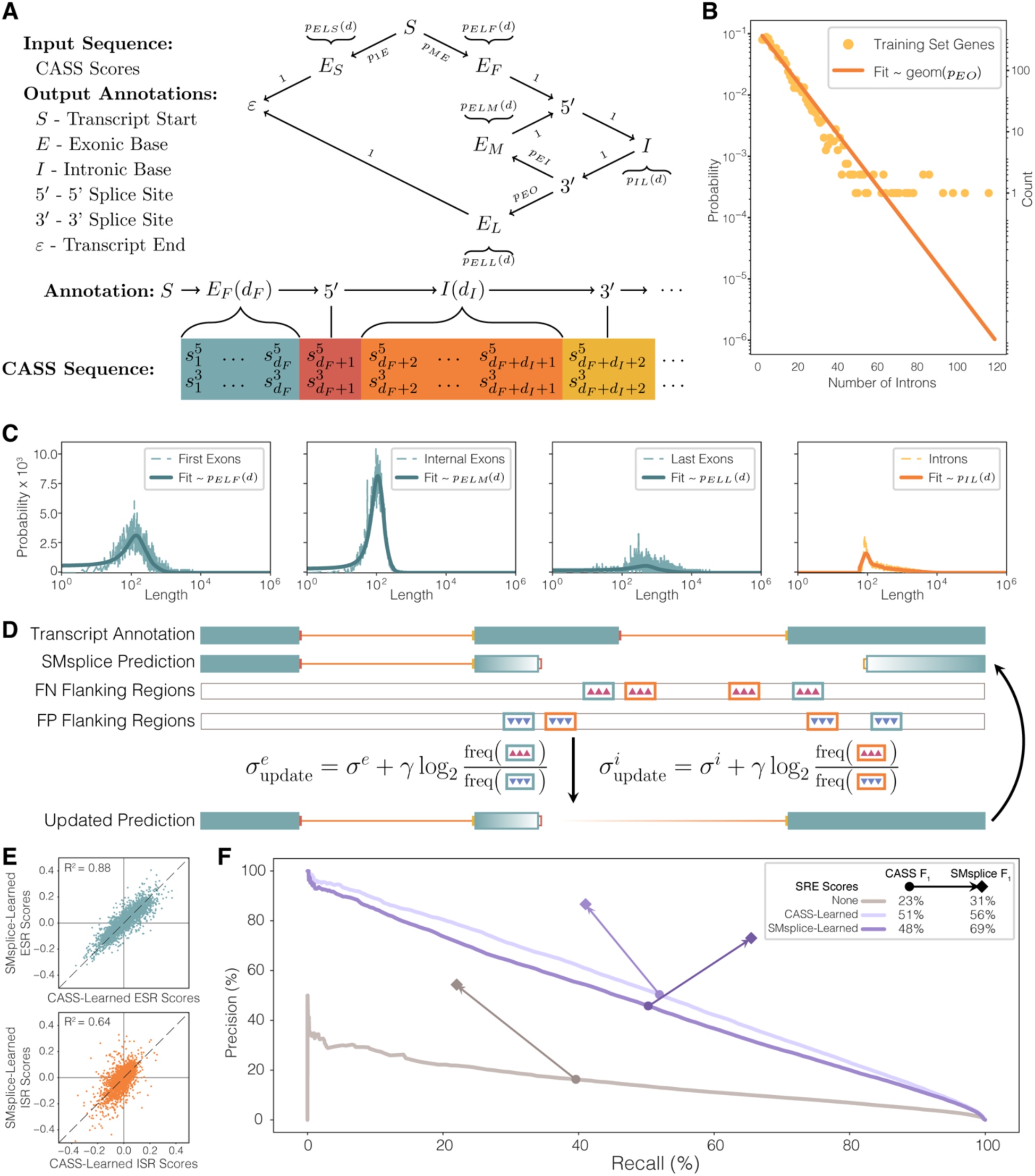
SMsplice Structure, Parameters and Performance in Human. A) In the upper diagram the labeled arrows represent the possible transitions with their associated probabilities, and the curly braces represent the association between states and their length distributions. Below, a possible parse of a sequence of CASS scores is shown, which would be determined from some RNA sequence of interest. B) Geometric fit to the empirical counts of genes with a certain number of introns in the training set. C) The empirical length distributions for the introns and first, internal (or middle), and last exons in the training set of genes along with the smoothed distributions used in the SMsplice model. D) Iterative learning of scores is performed as before, but using SMsplice predictions to define the false positive and false negative flanking regions. E) SMsplice-learned ESR scores correlate more with the CASS-learned ESR scores than their ISR counterparts. F) Precision-recall curves on the test set for the CASS framework predictions using different sets of SRE scores. The dots along the curves represent the cutoff learned on the training data to maximize the F_1_ score, and are connected by an arrow to a diamond marking the precision and recall for predictions made using the SMsplice model with the same SRE scores.

This structure reflects the observation that the number of introns per human gene fit approximately a geometric (exponential) distribution in the training set, with parameter *p*_*E*0_ = 0.093 (Figure 2B). We additionally characterized the length distributions of introns and different types of exons in human genes, which may reflect in part preferences involved in spliceosome assembly (De Conti et al., 2013) (Figure 2C). To model the distributions of first, internal, and last exons, we generated smoothed empirical length distributions on the training set for each type, substituting a geometric tail for each distribution to avoid issues with data sparsity at long lengths (Methods).

Our SMsplice model combines the above “structural” features (relating to exon/intron order/size), which we call the SMsplice structure, with the CASS scores. This model assumes that the exons and introns in a transcript are chosen in proportion to: 1) the strengths of the involved SS in local context, represented by (exponentiated versions of) the CASS scores; 2) the exon and intron length preferences represented in Figure 2C; and 3) the preferred number of introns represented by a geometric distribution (with parameter from Figure 2B). Together, these three features define the score for any “parse” (splicing pattern) *π* for the sequence of interest (‘seq’). If seq is of length *T*, an arbitrary parse *π* has *N* > 0 introns of lengths 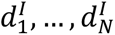, a first exon with length 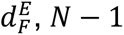 internal exons with lengths 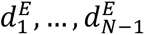, and a last exon with length 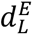, (with the sum of the lengths of all exons and introns being *T*). The SMsplice score for this parse, *SM*[seq, *π*], is then defined as:

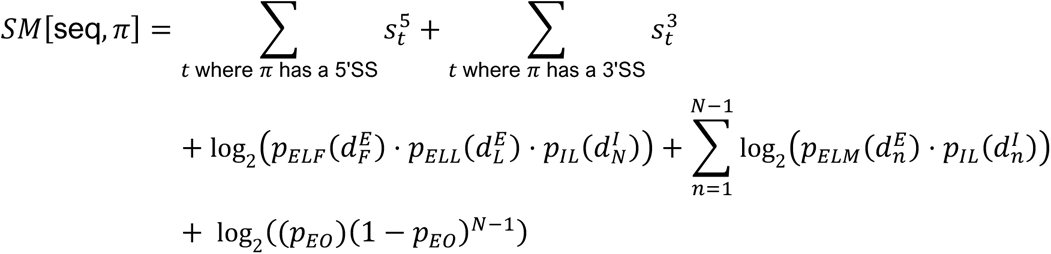

Here, the first line represents the strengths of the SS (in context, i.e. using CASS-style scores), the middle line represents length preferences for the involved exons and introns, and the last line represents the preference for the specific number of introns.

The predicted splicing pattern for the sequence, *π*^∗^, is the one which has the highest SMsplice score of all possible parses, which can be obtained by adapting the Viterbi algorithm for HSMMs (Methods) (Shun-Zheng & Kobayashi, 2003). While SMsplice is similar in structure to an HSMM, it is a discriminative rather than generative model, meaning that it discriminates different parses but cannot generate RNA sequences. Therefore, it is technically a semi-Markov conditional random field rather than an HSMM (Lafferty et al., 2001; Sarawagi & Cohen, 2004).

While we could, and do, use the previously determined CASS scores with SMsplice, we can also learn an additional set of SRE scores using SMsplice predictions. As before, we began by setting all SRE scores to zero, but now used the FP and FN from SMsplice predictions to update the scores rather than the CASS predictions (Figure 2D). We refer to the resulting SRE scores as SMsplice-learned. Comparing these scores with the CASS-learned SRE scores, we observed fairly strong correlation between the scores learned by both methods, more so for the ESR scores than the ISR scores (Figure 2E).

We then explored the predictions made using either set of SRE scores within either the CASS framework or SMsplice on a test set of genes (Figure 2F). This analysis showed that F_1_ performance improved with the addition of the SMsplice structure, regardless of the SRE scores considered, up to an increase of 21% over the CASS predictions. Furthermore, SMsplice had the highest performance with the SMsplice-learned SRE scores at 69%, substantially higher than with the CASS-learned scores. This improvement may result in part from the ability of SMsplice to exclude potential SS that are structurally incompatible with splicing that are considered by the CASS model; for example, a predicted 3’SS that occurs before the first putative 5’SS in a transcript (Fig. 1C). Additionally, SMsplice is more apt to predict SS pairs with typical exon/intron spacings and less apt to predict lone, high-scoring SS in introns than the CASS framework. Therefore, the false predictions considered during SMsplice-learning may be more enriched for regions considered by the splicing machinery whose exclusion is modulated by nearby SREs.

We further defined a “Local Score” (LS) for any intron or internal exon based on the associated terms in the *SM*[seq, *π*] expression. For an internal exon whose flanking 3’ and 5’SS occur at *t*1 and *t*2, respectively, resulting in an exon of length *d^E^* = *t*2 + 1 − *t*1 the LS is:

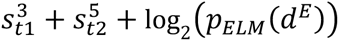

For an intron whose 5’ and 3’SS occur at *t*1 and *t*2, respectively, resulting in an intron of length *d*^1^, the LS is:

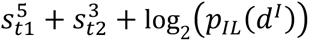

Applying these scores to predicted exons and introns for the human test set, we observed that exons with larger local scores were more likely to be predicted correctly, with a fairly smooth and nearly monotonic relationship, and similarly for introns (Supplemental Figure 1A).

### Application of SMsplice to other animals and a plant

Splicing is a fundamental process, thought to have been present in the last common ancestor of eukaryotes (Keren et al., 2010), and the assumptions about splicing underlying the CASS and SMsplice models are reasonable for a wide range of species. Furthermore, many protein families known to modulate splicing via the binding of SREs are conserved between animals and plants, though some have been lost especially in particular fungal lineages (Busch & Hertel, 2012). Therefore, it was of interest to explore the application of our models to genes from diverse organisms. To capture a range of evolutionary distances and address important model organisms, we selected mouse, zebrafish, fruit fly (*Drosophila melanogaster*), silkworm moth (*Bombyx mori*), and the model plant *Arabidopsis thaliana*. We devised separate training, validation, and test sets for each organism analogous to those used for human (Methods).

As the splice site motifs differ somewhat between these organisms, we trained a new organism-specific third-order MaxEnt model for each. These models were trained on all of the genes not in the organism’s test set, using the third-order constraints as was done with the updated human MaxEnt models (Methods). We then compared the classification performance of these organism-specific MaxEnt models to that of the original (human-trained) second-order MaxEnt model to assess the change in SS classification performance. We found that these new models provided improved discrimination compared with the original MaxEnt models for all 6 organisms, especially in *Arabidopsis* (Figure 3A).

**Figure 3:**
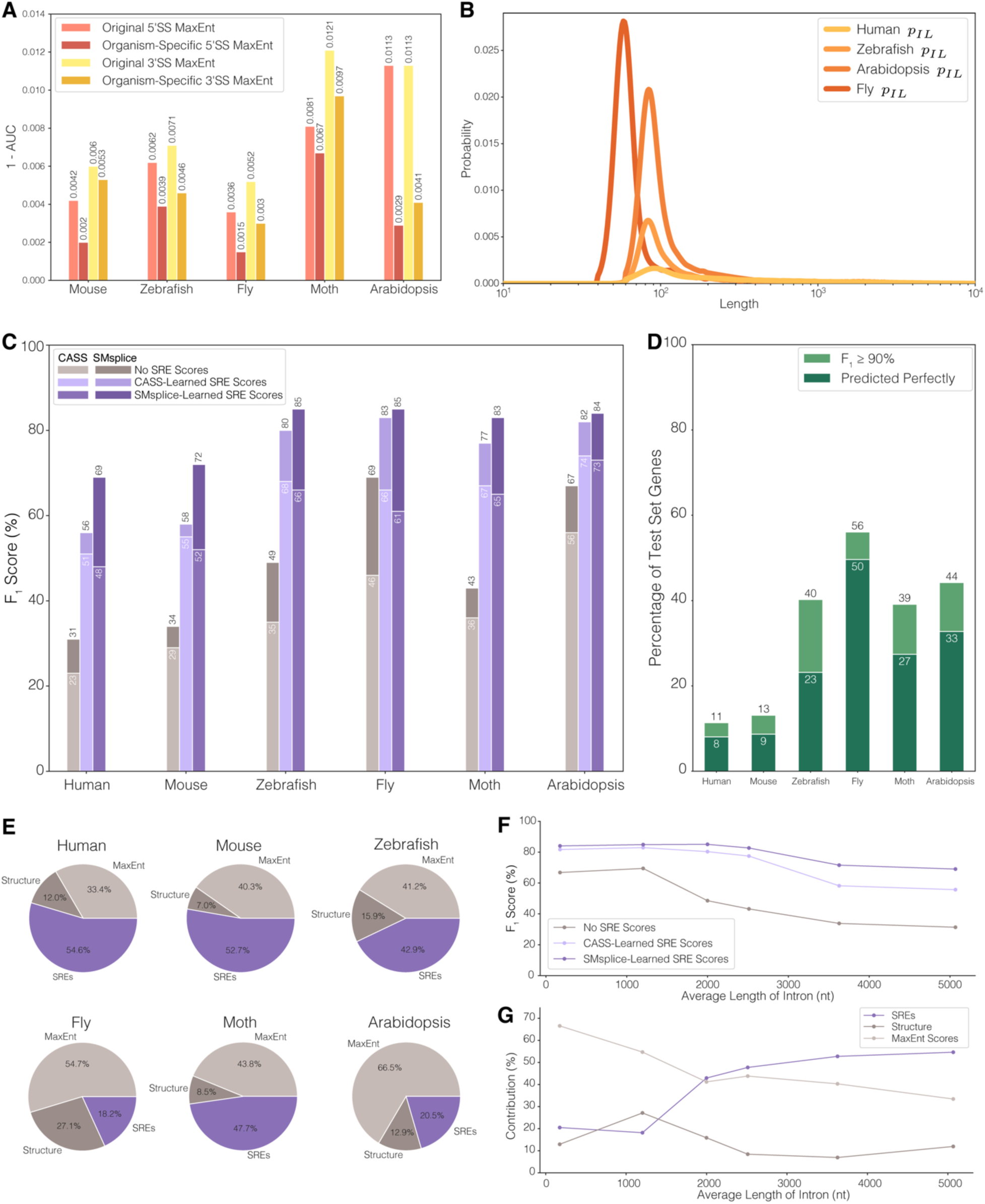
Application of SMsplice to six organisms show varied contributions of different features. A) Smaller 1-AUC values for the updated, organism-specific MaxEnt methods compared with the original human version show their improvement on classifying SS versus non-SS sequences. B) Intron length distribution fits used within the SMsplice model for human, zebrafish, *Arabidopsis*, and fly. C) Test set F_1_ performance for each organism with no SRE scores, CASS-learned SRE scores, and SMsplice-learned SRE scores within both the CASS framework and SMsplice model. D) Proportions of the test set predicted perfectly and with F_1_ score at least 90% for each organism. E) Pie chart breakdowns of the contributions to F_1_ performance of the organism-specific MaxEnt scores, the SMsplice structure, and the SMsplice-learned SRE scores. F) The F_1_ performance of the SMsplice model for each organism with no SRE scores, CASS-learned SRE scores, and SMsplice-learned SRE scores as a function of the average length of introns in the test set. G) The contributions to F_1_ performance of the organism-specific MaxEntScan scores, the SMsplice structure, and the SMsplice-learned SRE scores as a function of the average length of introns in the test set.

Using these new splice site models, we next learned organism-specific SREs using the CASS-learning and SMsplice-learning approaches. We also determined structural parameters for the SMsplice model, including exon and intron length distributions, for each organism (Methods, Figure 3B, Supplemental Figure 2) (Lim & Burge, 2001). Modeling exon and intron lengths was of particular interest as the length distributions for exons and introns are related to the prevalence of intron definition versus exon definition, with the latter being especially prevalent in mammals (De Conti et al., 2013; Keren et al., 2010).

When we applied these newly-determined parameters to the task of predicting splicing on test sets from the respective organisms, we observed several trends (Figure 3C). In every organism, use of CASS- or SMsplice-learned SRE scores improved performance substantially over SS scores alone. The SMsplice structure further improved performance, with best overall performance observed with SMsplice-learned SRE scores for every organism. While the highest F_1_ value for mouse (72%) was similar to the 69% seen in human, accuracy in the four other organisms were all substantially higher (83-85%). Some variation was seen in the relative performance of the different score/framework combinations, with a larger gap between SMsplice- and CASS-learned SRE scores for mammals than other organisms, perhaps reflecting the longer distances involved in mammalian splicing. The local scores of exons and introns in these organisms showed the same positive association with accuracy as was observed for human (Supplemental Figure 1), indicating that this score can be used to distinguish predictions of higher and lower confidence. Thus, our framework and learning approach generalize well to other animal and plant species, enabling a variety of comparative investigations.

We also assessed performance at the level of individual genes, again using the F_1_ measure. For most organisms, the median F_1_ accuracy for individual genes in the test set was close to or slightly above the overall F_1_ value (Supplemental Figure 3). The exception was fly, whose median test gene had an F_1_ of 95%, reflecting that nearly >49% of the test genes were predicted with perfect accuracy. The proportion of perfectly predicted genes in other organisms varied from 8% (human) to 33% (*Arabidopsis*) (Figure 3D). A representative splicing prediction for each organism – with F_1_ value within 1% of the median value across genes – is visualized in Supplemental Figure 4. Examination of these visualizations reveal several common features of splicing, including the high density of decoy SS, the somewhat lower density of locations with high CASS scores, and a complex landscape of SRE scores. Many genes in *Arabidopsis* differed from those in other organisms in that there was often a visually apparent shift in ISR and/or ESR scores at or very near exon/intron and intron/exon boundaries that often persisted throughout the succeeding exon or intron (Supplemental Figure 4).

### Splicing in organisms with longer introns is more dependent on SREs

One of our fundamental goals was to understand the relative contributions of different features to splice site recognition. The CASS framework was designed to mirror the concept of SRE regulation of SS motifs, and these two components can be separated into the SRE scores and MaxEnt scores, respectively. Then, the SMsplice structure additionally includes the structural constraints of the spliceosome. So, by examining their effects within the model, we might gain some insight into how important these different types of splicing information are in these organisms. To do so, we considered the change in performance provided by the use of the SMsplice structure in the absence of SREs, and then the further benefit that adding the SMsplice-learned SRE scores granted (Figure 3E). From this, we noted that there was substantial variety in the proportions across these organisms. Human and mouse, for instance, saw the majority of their performance come from the SRE scores, while *Arabidopsis*, with the shortest mean intron length, had the most reliance on core SS motif scores. The largest structural contribution was seen in fly, which has a large proportion of introns within a very narrow length range of 40-80 nt (Pai et al., 2017).

Intron length varied substantially more than exon length across the organisms considered, and determines to a large degree the number of decoy SS per transcript (Black, 1995; Z. Wang et al., 2004). Noting this, we explored the relationship between intron length, performance, and the contributions of different features. In general, SMsplice performance declined as average intron length increased (Figure 3F). However, the decline was much shallower when SREs were included in the model. Consistent with this observation, we found that the relative contribution of the SS motifs to performance decreased with average intron length, while the relative contribution of SREs increased dramatically (Figure 3G). These trends suggest that, in lineages where introns lengthen, dependence on the presence of SREs becomes stronger.

### Learned SRE Scores Distinguish Real and Decoy SS

To further explore the idea of distinguishing real and decoy SS via SRE regulation within the model, we began by determining a suitable set of decoy SS. To create such a set, for each SS in the canonical training set that flanked an internal exon, we selected a non-SS base whose associated SS score was within half a bit of the score of the true SS (Methods). Thus, we were able to create a decoy set of similar size and score distribution to the set of real SS for each organism. We considered the distribution of SMsplice-learned SRE scores relative to the selected decoy 5’SS and 3’SS, as well as real 5’SS and 3’SS. In our models, 5’SS scores are affected by upstream ESRs and downstream ISRs, and 3’SS are impacted by upstream ISRs and downstream ESRs. For each position in the flanking regions, we considered the average ESR or ISR score (Figure 4A). Dividing the ESRs into ESEs (if score > 0) or ESSs (if score < 0) and similarly dividing ISRs into ISEs and ISSs, we separately tallied the averages of each of these categories of SREs.

**Figure 4:**
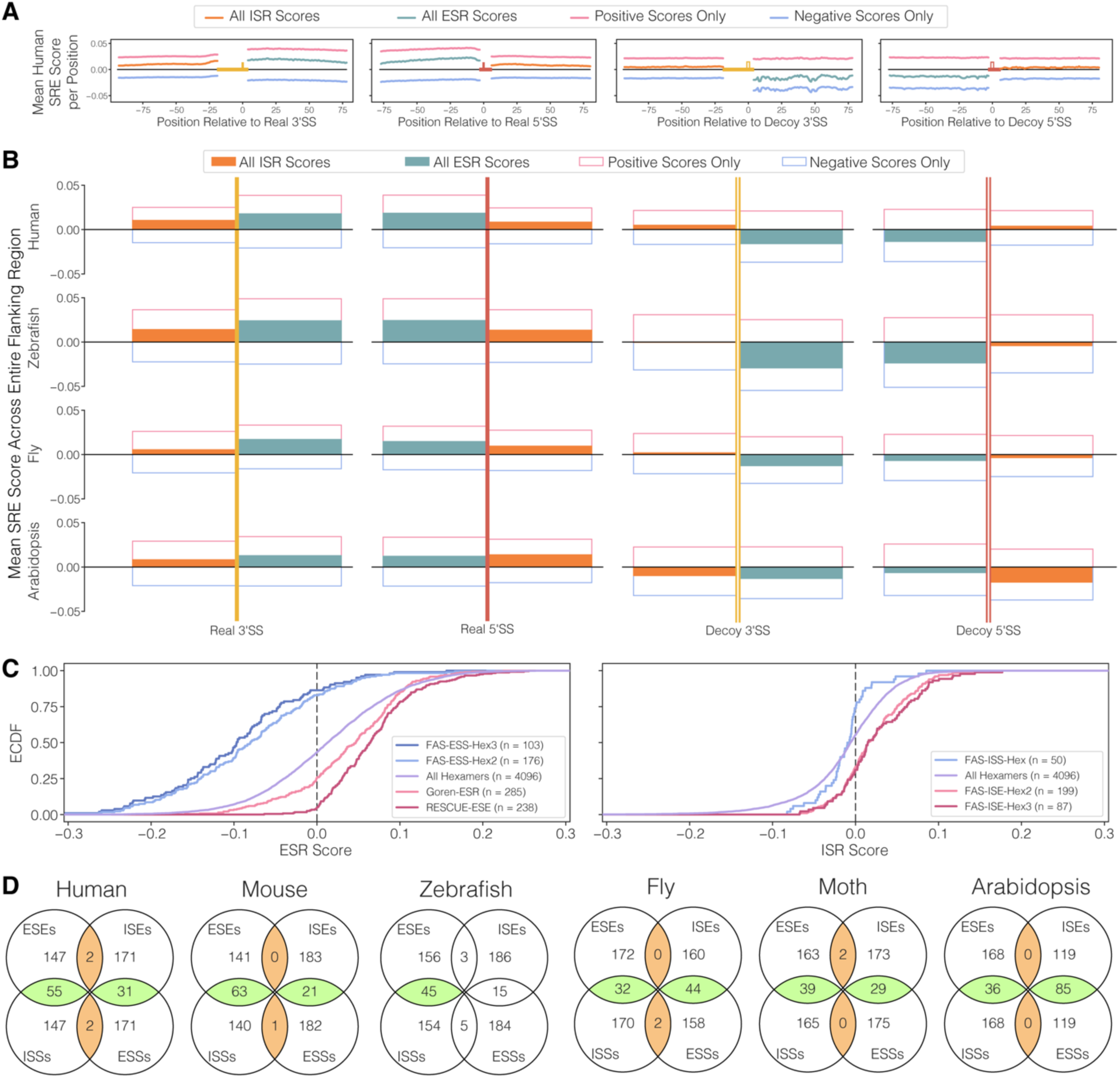
SMSplice-Learned SRE scores have features associated with known SREs. A) The relevant SMsplice-learned ESR and ISR scores, along with their positive and negative components, for each position in the flanking regions, averaged separately across all real and decoy 5’SS and 3’SS in the human training set. B) For human, the average of the scores shown in 4A across the associated regions. For zebrafish, fly, and Arabidopsis, the equivalent values on their training sets. C) Plots of the ECDFs for the human SMsplice-learned ESR and ISR values of all hexamers along with subsets of hexamers reported in the literature as particular types of splicing regulators in human. D) Agreement between the top and bottom 5% hexamers based on their SMsplice-learned ESR and ISR scores. Overlaps that are significantly under-represented are colored in brown, while overlaps that are significantly over-represented are colored in green.

In human, the average positional ESR and ISR scores flanking real SS were relatively smooth, as were the average ESE, ESS, ISE, and ISS values, suggesting that SRE information is on average fairly evenly distributed (Figure 4A). We did observe a modest increase in ESE scores at both exon boundaries, as has been observed previously (Cáceres & Hurst, 2013). Similar slightly sloping ESE distributions were observed in mouse, zebrafish, and *Arabidopsis* (Supplemental Figure 5A). ESS scores were also fairly smooth, with slightly increased magnitude further from both 5’ and 3’SS. The average SRE scores flanking decoy splice sites were also fairly smooth, with the exception of ESSs near decoy 3’SS, which showed a visually distinctive pattern of peaks and valleys, with a less pronounced pattern observed near decoy 5’SS. Suspecting that a common repetitive element might be involved, we observed that 18% of decoy 3’SS and 14% of decoy 5’SS overlapped SINE elements of the Alu class, often at particular locations (Keren et al., 2010). Examining the SRE distributions in the Alu vs non-Alu decoys, we found that the decoy SS in Alus had a more exaggerated version of the overall pattern of peaks/valleys, indicating that Alu decoys are driving this pattern (Supplemental Figure 5B). In some other organisms (e.g., mouse, moth), similar bumpiness in ESS distributions was observed (Supplemental Figure 5A), likely for similar reasons.

To summarize the impacts of different SRE score categories on SS prediction, we also considered the average of the relevant scores for the entire flanking regions (Figure 4B, Supplemental Figure 5C). For every organism, there was positive net score for real versus decoy 5’ and 3’SS in both the exonic and intronic flanking regions, as expected. Furthermore, the average ESR score was of greater magnitude than the average ISR score for real 5’ and 3’SS in every case, except the 5’SS of *Arabidopsis*. For the decoys, we observed a negative average ESR score on the exonic side for both splice site types in all organisms. ISR scores on the intronic side of decoy SS were more variable in sign, with smaller magnitude than the corresponding ESR scores in all cases, except for decoy 5’SS in *Arabidopsis*.

The patterns we observed for the SRE scores in regions flanking decoy SS suggested that these regions might be useful in SRE score learning. To explore this idea, we scored each hexamer according to its frequencies of appearance in the CASS-relevant regions flanking the real and decoy SS in the training set, and found that these “real versus decoy” scores correlated strongly with the SMsplice-learned SRE scores (R^2^ values between 0.34 and 0.79, Supplemental Figure 6A, Methods). Furthermore, when appropriately weighted, these real versus decoy scores provided useful seeds to SMsplice-learning, providing a small but consistent improvement in validation performance of ∼1% over no seeding (Supplemental Figure 6B, Methods). Going forward, we used the scores learned from these seeds as our SMsplice-learned SRE scores. As observed for the CASS scores, the new ESR scores tended to have greater magnitudes than the ISR scores in human (Figure 1E, Supplemental Figure 6C). For the non-mammals, ESE scores tended to exceed ISE scores, but ISSs often had greater average magnitudes than ESSs.

To ask how the SRE scores learned by our SMsplice model relate to known sets of SREs, we gathered several groups of hexamers determined to function as ESSs, ESEs, ISSs and ISEs using splicing reporter-based screens or validations (Fairbrother et al., 2002; Goren et al., 2006; Y. Wang et al., 2012, 2013; Z. Wang et al., 2004). We compared the empirical distributions of SRE scores associated with these different groups of hexamers to the relevant overall empirical distribution of SMsplice-learned SRE scores (Figure 4C). All of the ESR groups were significantly different from the overall ESR score distribution (*p* < 10^-3^, two-sample KS test), and the ISR groups were also significantly different from the overall ISR score distribution (*p* < 0.01, two-sample KS test). Furthermore, RESCUE-ESEs predominantly had positive ESR scores (226 out of 238, 95%), and ESRs from Goren et al were also strongly enriched for positive scores (both *p* < 10^-3^, one-sided binomial test). Furthermore, both sets of FAS-ESS hexamers were strongly over-represented for negative ESR scores (89 out of 103, or 86% for the FAS-hex3 set) (*p* < 10^-14^for both sets, one-sided binomial test). These trends were also observed for intronic elements, with FAS-ISEs over-represented for hexamers with positive ISR scores, and FAS-ISSs over-represented for negative ISR scores (*p* < 0.01, one-sided binomial test). These observations support that positive/negative SRE scores learned by our model are predictive of splicing enhancing/silencing splicing activity, respectively.

It has been previously observed that human SREs have a pattern of agreement where ESEs and ISSs tend to overlap as do ISEs and ESSs, likely reflecting common mechanisms of the associated RNA-binding splicing factors (Y. Wang et al., 2012, 2013). To explore whether our SMsplice-learned SRE scores recapitulated this pattern, we took the hexamers with most extreme 5% ESR and ISR scores. The top 5% scoring hexamers we considered our enhancers (ESEs and ISEs), and the bottom 5% scoring hexamers we considered our silencers (ESSs and ISSs). While 5% is an arbitrary threshold, it yields sets of 204 hexamers, similar in size to the SRE sets from the literature discussed above. Examining the hexamers in common between these four groups of 204 hexamers, we saw that there was indeed a great deal more overlap between the ESEs and ISSs, and the ISEs and ESSs than between the other options of overlap. Tests for over- and under-representation found that all of these overlaps were significant in the expected direction (*p* < 0.05, one-sided binomial test, Bonferroni-corrected) aside from the comparisons involving zebrafish ISEs and ESSs (Figure 4D). These observations provide further support that strongly-scoring hexamers in our model possess properties expected of the associated SRE class.

### Clustering human hexamers recovers known splicing RBP motifs and significant splicing regulators

Known SREs predominantly function by recruitment of splicing regulatory factors that bind motifs of approximately 3-7 nt in length (Ray et al., 2013). To ask whether the SMsplice-learned SRE scores correspond to known splicing regulatory factor motifs, we clustered hexamers from the sets of 5% most extreme positive and negative ESR and ISR scores considered above. To this end, we clustered the component hexamers of each set by sequence similarity and aligned them to create a position weight matrix (PWM) for each of the resulting clusters (Supplemental Figures 7-10, Methods). The ESS clusters were notably poor in cytosine, consistent with a previous cell-based screen for ESSs (Z. Wang et al., 2004). For each of the clusters obtained from human SRE hexamers, we identified the best-matching motif from the set of RNAcompete in vitro RBP-binding motifs (Ray et al., 2013) using a simple permutation approach to assess which matches had significant similarity (Methods).

From this analysis, we identified an RNAcompete match that was significantly similar for the majority of human SRE clusters (Figure 5A, 5B, Supplemental Figure 11). The matching was particularly strong for the ESE and ESR clusters, where 15 out of 20 had a significant RNAcompete match. For the ESE clusters, matching occurred to motifs for known splicing factors including RBM45 and SRSF3, as well as other RBPs with homologs involved in splicing (PCBP4) or other roles in RNA metabolism (e.g., Star-PAP). For the ESS clusters, matching occurred to several known splicing factors, including HNRNPA1L2, HNRNPA3, HNRNPDL, the HNRNPF/H family, the MSI family (whose motif resembles that of HNRNPA0), and QKI.

**Figure 5:**
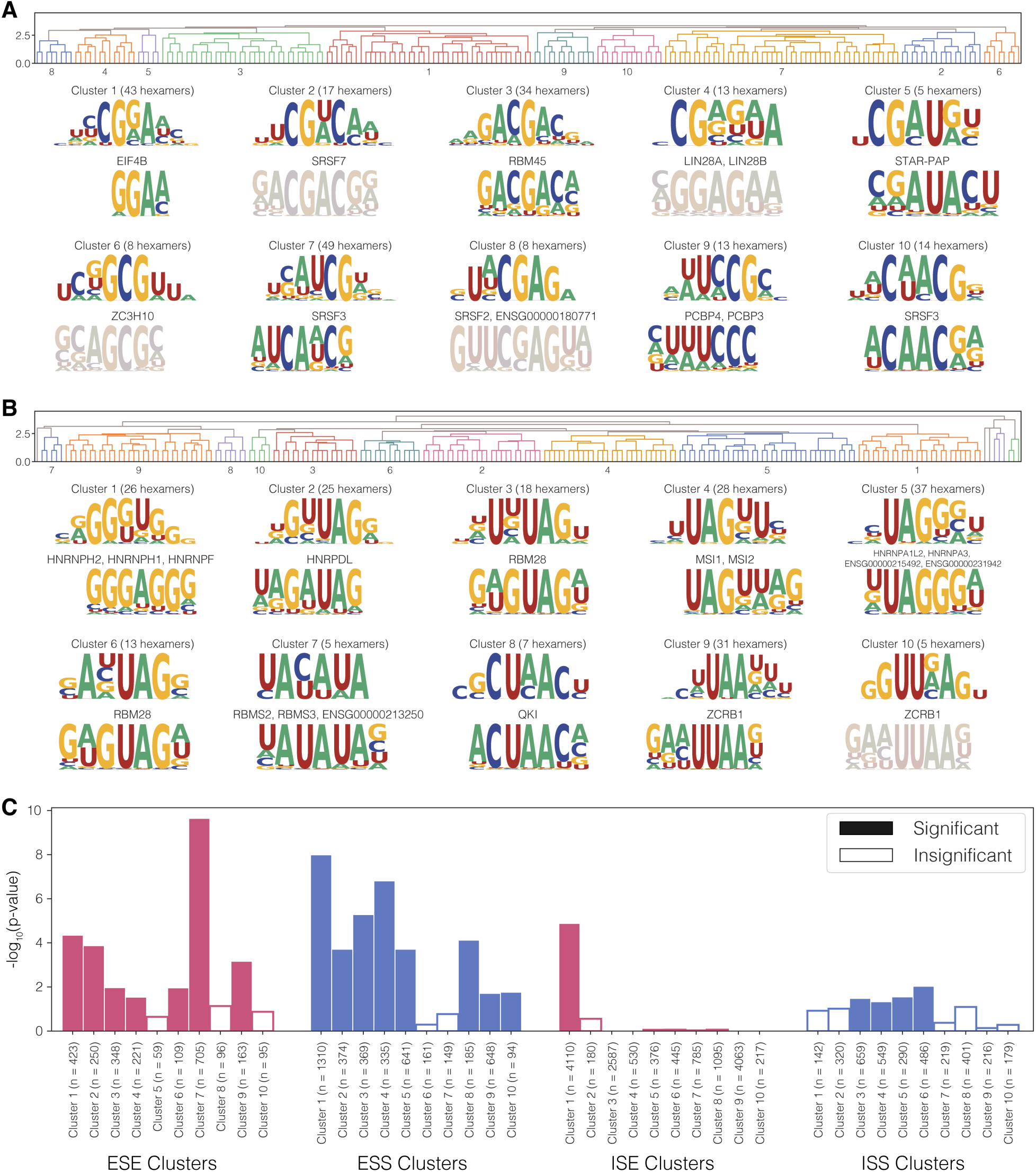
Human SRE cluster motifs often match known RBP binding sites. A) Dendrogram for clustering hexamers with the 5% most positive ESR scores in human, as well as the resulting aligned clusters and best RNAcompete matches below those clusters. Desaturation indicates that the best match did not satisfy our threshold. B) Dendrogram for clustering hexamers with the 5% most negative ESR scores in human, as in A). C) For each of the clusters made from the sets of the most extreme scoring hexamers, the support for that cluster being more associated with changes in splicing outcomes. The number of events available for each cluster is indicated, as well as whether the association is deemed significant.

Thus, most of the ESR motifs obtained by this clustering matched known splicing factors motifs. ESS motifs primarily matched hnRNP motifs, most of which are known to repress splicing when bound to exonic locations, and two ESE clusters matched SR proteins, which are known to activate splicing from exonic locations (Busch & Hertel, 2012). Many of the ISR clusters also matched known splicing factor motifs, including ISS clusters that matched RBM42, SRSF3, and SRSF7 motifs, and an ISE cluster that matched the QKI motif (which also resembles the branch point and SF1 motifs). Together, these observations suggest that our score learning and clustering approach identifies authentic SRE motifs without prior knowledge of RBPs guiding the process. Clusters that fail to match RNAcompete motifs may represent known splicing factors that have not yet been characterized in vitro, motifs for unknown splicing factors, or artifacts of the algorithm or clustering process.

We further wanted to assess the splicing regulatory activity of our clusters to see if the hexamers that we identified as high-scoring were more associated with changes in splicing outcomes. To do this, we used fine-mapped splicing quantitative trait locus (sQTL) data from the Genotype-Tissue Expression (GTEx) database to ask whether variants that disrupted hexamers in each cluster were more likely to be causal relative to control hexamers (Methods). This analysis supported the regulatory activity of seven ESE clusters, eight ESS clusters, one ISE cluster, and four ISS clusters (Figure 5C). These supported clusters included ESS cluster 10, and ESE clusters 2, 4, and 6, which had no sufficiently strong matches within the RNAcompete set. This observation suggests that the hexamers identified by our model could represent SREs that are as yet unassociated with known RBPs. The greater number of exonic clusters than intronic clusters that reached significance could reflect stronger activity of ESRs than ISRs, or better modeling of ESR than ISR elements by our model.

### Similar regulatory motifs identified across organisms allows for generalization of SMsplice model

Key splicing proteins of the SR and hnRNP classes are conserved from animals to plants (Busch & Hertel, 2012), suggesting that similar motifs may exert similar effects on splicing. However, intron-containing genes are often spliced in a different way when moved between mammals and fish (Yeo et al., 2004), and potentially even more so across greater evolutionary distances. Examining the clusters for each SRE class in the non-human organisms (Supplemental Figures 7-10), we observed many similar motifs within each class.

We repeated our motif comparison analysis for each examined organism, matching the clusters to RNAcompete RBP motifs associated with the relevant organism (Supplemental Figures 12-16). These comparisons identified a number of RBP matches in each organism, generally to very similar classes of proteins as observed in human, with several ESEs matching SR proteins, ESSs often matching hnRNPs or the MSI family, and QKI family proteins appearing as ISE matches. These observations suggest that clusters identified by our algorithm in other species often correspond to motifs bound by splicing factors, many of which are conserved.

To explore the extent of similarity of ESS clusters across organisms, we clustered all the ESS PWMs across species, using the same distance measure used to compare our PWMs with the RNAcompete PWMs (Figure 6A). This comparison identified several motifs, including UAG- and UAA-containing motifs, that were present in all six organisms, suggesting that these could represent ancient ESSs present in the last common ancestor (LCA) of animals and plants. Other motifs, including poly-G motifs, were present in all metazoans but were not observed in *Arabidopsis*. Some of the more CG-rich ESSs appeared more lineage-specific. For instance, a CGCG motif only identified as an ESS in zebrafish resembled motifs identified as ISSs in other organisms (Supplemental Fig 16).

**Figure 6:**
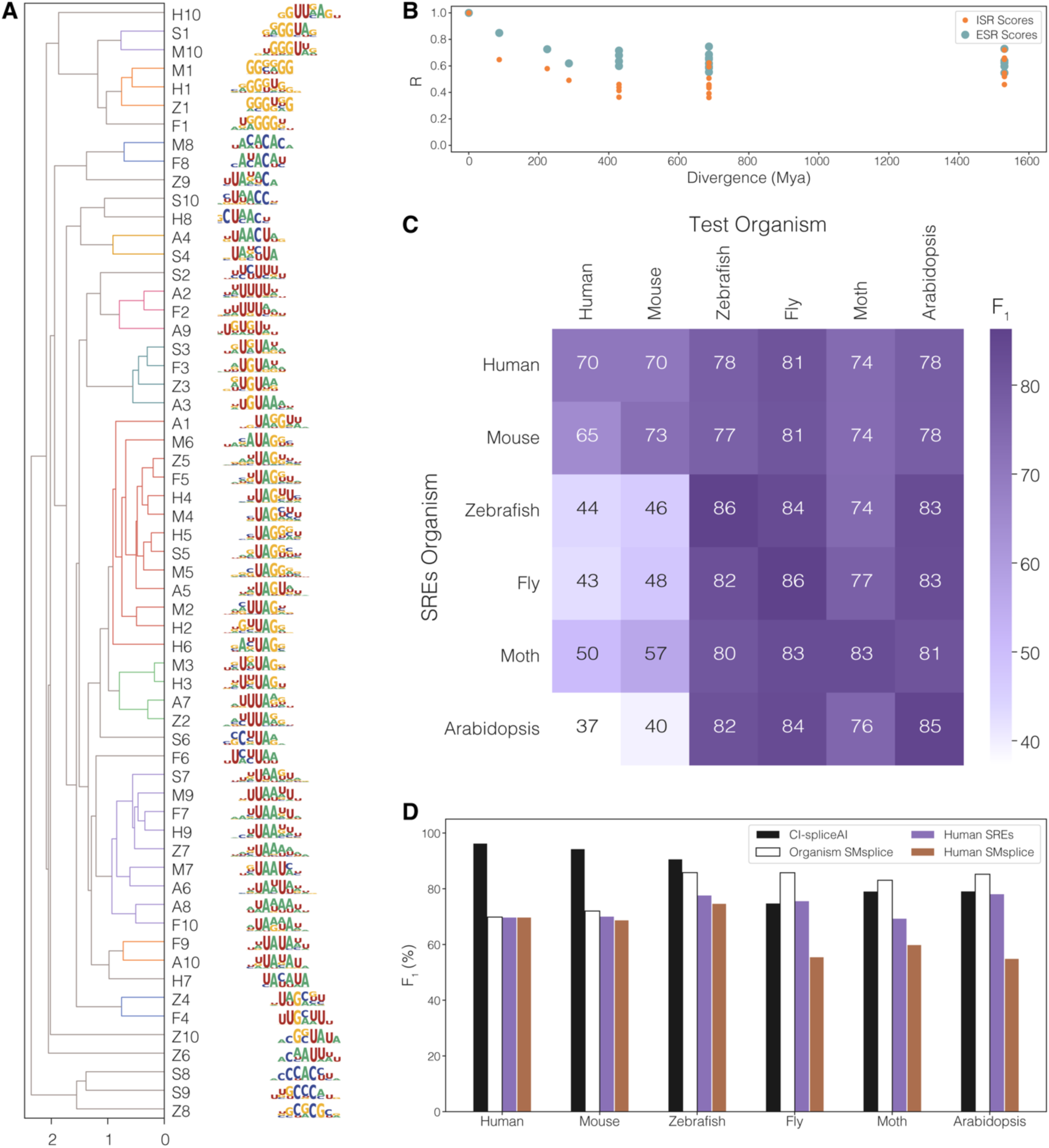
Comparing SRE motifs and splicing models across organisms. A) All ESS clusters for all organisms clustered by motif similarity. Motifs are labeled using the first letter of the organism, using “S” for silkworm moth, followed by the number of the ESS cluster in Supplemental Figure 8. B) Correlation between ESR scores and ISR scores for each pair of organisms (as well as fugu, data otherwise not shown) as a function of their evolutionary divergence. C) F_1_ performance on each organism using weighted SREs from each other organism. D) F_1_ performance on each organism for CI-SpliceAI, the SMsplice model with all organism-specific parameters, the SMsplice model with SRE scores from human, but otherwise all organism-specific parameters, and the human SMsplice model.

We performed similar clustering analyses of ESEs, ISEs and ISSs across the six organisms (Supplemental Fig 16). For each class, we observed clear clusters, with some spanning all six organisms or the five metazoans. For example, we observed a cluster of ESE motifs related to “CYACG” (Y = C or U) in all six organisms, a cluster of ISE motifs containing “UAAC” in five of six organisms (all but moth), and an ISS motif cluster containing CGG and/or GGA, also present in all organisms except moth. Other clusters were more lineage-restricted, including a poly-G ISE motif in vertebrates and moth, and a purine-rich ISS motif in the four non-mammals. These observations suggest the presence of both conserved and lineage-specific SRE motifs across the species considered.

As the motifs were derived from subsets of hexamers with the most extreme scores, we wondered whether the full sets of SMsplice-learned SRE scores would also exhibit conservation across species. To compare ESR and ISR conservation, we calculated the correlation of scores across ESR and ISR scores as a function of evolutionary divergence time (Figure 6B) (Kumar et al., 2022). We observed a positive correlation of ESR and ISR scores across all pairs of organisms, further supporting that some SREs have maintained similar activity across > 1.5 B years of evolution. While very close organisms had more similar ESR scores, the similarity plateaued at a moderate correlation at evolutionary distances of ∼400 Mya and beyond. Notably, ESRs tended to be somewhat more conserved than ISRs, consistent with the high conservation of ESS and ESE motifs observed above (Cáceres & Hurst, 2013), and the fact that the protein-coding function of exons provides a strong constraint on their compositional drift that is not present in introns.

We were curious whether the SRE similarity we observed above was strong enough that one organism’s SMsplice-learned SRE scores could be used effectively to predict splicing in the genes of another organism when paired with core SS motif scores and structural parameters of the other organism. Applying SMsplice in this manner with all possible combinations of SREs/genes (Methods), we observed that SREs from the non-mammalian species performed well on other non-mammals but poorly on the mammals. This finding may relate to our observation that mammals derive far more information from SREs than other organisms (Figure 3D) and have some lineage-restricted SRE motifs (Figure 6A, Supplemental Figure 17). Perhaps the intrinsic difficulty of identifying SS using local information in longer mammalian transcripts necessitates more species-optimized SRE scores than in other lineages. On the other hand, the mammalian-trained SREs performed reasonably well on all organisms, always outperforming the organism-specific SMsplice model without SREs (Figure 3C), suggesting a reduced prominence of lineage-specific SREs in non-mammals.

The variations in the other components of the SMsplice model between organisms play a substantial role in the ability to generalize across species. For instance, using the human SMsplice-learned SRE scores within an organism’s SMsplice model always outperformed the fully human-specific model (Figure 6D). In every case, the organism-specific SMsplice model had the best performance, as expected. As a point of comparison, we also applied the neural net model CI-SpliceAI to the same test sets (Strauch et al., 2022). This black box model was trained on human sequences and so the most direct comparison is to the SMsplice model with fully human parameters. Comparing performance, CI-SpliceAI had substantially higher performance on human and mouse genes than our human SMsplice model, but did not generalize as well to non-vertebrates. Furthermore, our organism-specific SMsplice models outperformed CI-SpliceAI on all non-vertebrate organisms. As CI-SpliceAI is a black-box model, we cannot consider extracting parameters related to SREs, for example, as we could for the SMsplice model, so it was not feasible to mix and match CI-SpliceAI with SMsplice. One of the advantages in structuring a model in an explicit manner as we have done is that it allows for the exploration and interpretation of the different parameters of the model as shown in Figures 6C and 6D.

## Discussion

A few simple assumptions about features important for splicing underly SMsplice, principally that SS recognition depends on a core motif modulated by nearby SREs, and that SS are recognized in pairs with favored lengths for both exons and introns. The model depends on just three types of parameters: the triplet-based core SS motif model, structural parameters capturing preferences for particular exon and intron lengths and numbers, and scores of hexamers as exonic and intronic SREs. The core SS motif and structural parameters are directly estimated from frequencies in the training data, with no adjustments or “learning” involved, thus generating a scaffold for SS recognition in the absence of SRFs. Scores of hexamers as exonic and intronic SREs were then learned based on their ability to predict splicing patterns within this scaffold, yielding SRE clusters that were predictive of the splicing effects of genetic variants.

The organism-specific SMsplice models enabled us to explore many facets of splicing across species. For example, we found that organisms with longer introns have greater reliance on SRE scores, and that *Drosophila* had the greatest reliance on the structural parameters, likely related to strong preferences for a narrow range of short intron lengths in flies (Talerico & Berget, 1994). Additionally, we generally observed that ESR scores had greater prominence than ISR scores except in *Arabidopsis* where ESRs and ISRs appeared comparably important, consistent with early studies showing that U-rich sequences distributed throughout plant introns are important in intron recognition (Lorkoví et al., 2000).

In human, where SREs and the effects of mutations on splicing outcomes have been extensively studied, we observed significant agreement between our learned parameters and experimentally validated sets of SREs (Fig. 4C), and the top and bottom 5% of ESR and ISR hexamers yield clusters that match well to binding motifs of many known SRFs (Fig. 5, Supp. Figs 11-16). In other organisms, agreement with the somewhat sparser sets of known RBP binding motifs available was also observed, with additional clusters representing candidate novel SREs in each lineage. Unlike qualitative catalogs of SREs identified previously, our approach learns a score for each hexamer, distinguishing stronger and weaker ESEs, ESSs, ISEs and ISSs. This feature could be particularly useful for predicting the most essential SREs governing splicing of a given exon, intron or splice site, with potential applications for interpretation of genetic variants or design of splice-switching antisense oligonucleotides (ASOs) (Havens & Hastings, 2016).

In addition to the full SMsplice model, the new core SS motif models and CASS framework may have useful applications in assessing the strength of a SS in local context. These models can be applied even on short sequences of a few dozen bases (for SS motifs) or one to two hundred bases (for CASS), where neural models typically struggle. For example, CASS scores might find use in design or troubleshooting of splicing in minigenes, or in synthetic biology applications. The SMsplice structure, which normally uses our new MaxEnt SS scores, could potentially also be applied to SS scores derived in other ways (Cheng et al., 2019) because of its modular design.

While we have learned model parameters for several organisms, a predictive tool to study splicing might be useful in many other organisms. In some cases, using the pre-trained model for the evolutionarily closest organism may suffice, e.g., using the zebrafish model to predict splicing in other fish species. However, given sufficient high-quality annotated gene data, organism-specific parameters can be generated, using the provided code. New structural parameters and MaxEnt models are straightforward to derive, while SRE score learning is more computationally intensive. Our experiments have shown reasonable generalization of SRE scores within mammals and within the non-mammals studied, while structural features vary substantially across lineages.

Therefore, simply deriving new SS models and structural parameters with pre-trained SRE scores may provide most of the benefits of a fully organism-specific splicing model with relatively modest effort.

### Limitations of this work

The SMsplice framework was designed with typical animal and plant splicing systems in mind, and is likely less suitable for modeling splicing in fungi, where the BPS motif is of much greater importance (Kupfer et al., 2004). Furthermore, our training and testing procedures are dependent on the accuracy of the underlying genomic sequences and annotations, potentially limiting application to organisms where annotations are less accurate. Additionally, we have considered canonical annotations, which represent predominantly constitutive exons, and have not explicitly explored alternative splicing in this study. Another limitation is the fact that, while we have attempted to construct and learn the SRE scores in such a way that they reflect splicing regulatory activity (e.g., emphasizing hexamers within ∼80 nt of the SS), it is inherently challenging to isolate such activity from biases in sequence composition of exons and introns that reflect other facets of gene expression, such as protein coding, mRNA export or stability, and/or processes of DNA mutation and repair. While we have shown that hexamers with the most extreme SRE scores are strongly associated with SRF binding and splicing activity, weaker-scoring SRE hexamers may be less enriched for splicing regulatory activity. To enhance tractability and interpretability, we have also made the strong assumption that SREs act only over short distances and interact only additively. This assumption ignores potential long-range activities and synergistic or antagonistic relationships that undoubtedly exist in some cases. These limitations represent worthwhile challenges for the next generation interpretable splicing models.

## Acknowledgements

We thank Sean Eddy and Gene-Wei Li (members of K. M.’s thesis committee), as well as Armando Solar-Lezama, Phillip Sharp, Osbert Bastani, Kavi Gupta, and Chenxi Yang for helpful discussions, members of the Burge lab for helpful comments, and Hannah Jacobs for assistance with the fine-mapping data and analysis. This work was supported by grants from the NSF (PI: A. Solar-Lezama), the NIH (C.B.B.) and a Computational Sciences Graduate Fellowship from the Department of Energy (to K. M.).

## Methods

### Software and Data Availability

The code for SMsplice, and code for learning new 3^rd^-order MaxEnt splice site models and SRE scores is provided on Github (http://github.com/kmccue/smsplice), as are the datasets used in training and testing that are described below.

### Datasets

The datasets used for analyses of human splicing were based on the canoncial_dataset.txt used in training and testing of SpliceAI, in the hg19/GRCh37 genome version (Jaganathan et al., 2019). We excluded from consideration any genes whose sequence contained ’Ns’ or non-standard bases, or whose total length was less than 100 bases or whose shortest annotated intron was less than 25 bases (unrealistically short, suggesting an annotation error). From the genes which would not be in the test set we defined a validation set by randomly selecting 1,000 non-paralogous genes. We then randomly selected 4,000 genes from the remaining genes which would not be in the test set to act as the training set. An additional set of 1,000 randomly selected genes which would not be in the test set was chosen as our SRE weight learning set. For practical (speed) reasons, we excluded genes longer than 200 kb from the validation and weight learning sets above. We used the same test set as SpliceAI, applying the filters as above, which removed ∼1% of genes, resulting in a set of 1,629 genes. Aside from the MaxEntScan training described below, these sets were used throughout the paper.

For each non-human organism, to focus on the most highly used and reliable splice sites, we created a canonical dataset containing a canonical splicing pattern for each protein-coding, intron-containing transcript that met the filtering criteria used for human genes. We also created an “all-SS” dataset containing the full complement of splice sites present within the respective canonical transcript, analogous to the gtex_dataset.txt from SpliceAI. Training, validation and test sets were derived from the canonical dataset. The all-SS datasets were used for MaxEnt training and the real versus decoy splice site analyses.

For mouse, we used the knownCanonical table for the GRCm38/mm10 assembly to create the mouse canonical dataset, excluding the handful of genes with more than one annotation, and used the knownGene table to create the all-SS mouse dataset. We downloaded paralog annotations from Ensembl’s BioMart. The training, SRE weight learning, and validation sets were also selected from the genes not used in the test set as described for human. Nonparalogous genes on chromosomes 1, 3, 5, 7, and 9 were used to form the test set, resulting in a set of 1,212 genes.

For zebrafish, splicing annotations for GRCz11 were downloaded from Ensembl’s BioMart along with paralog status and APPRIS annotations (Rodriguez et al., 2013). To select the canonical transcript we filtered for APPRIS principal transcripts for each gene, and selected the longest transcript for genes with multiple APPRIS principal transcripts. Genes without a unique longest APPRIS principal transcript were discarded. The training, SRE weight learning, and validation sets were selected as in human from genes that were not used in the test set, which was made of nonparalogous genes on chromosomes 1, 3, 5, 7, and 9, resulting in a set of 825 genes.

For *Drosophila*, we downloaded splicing annotations, APPRIS annotations, and paralog status for dm6 from Ensembl’s BioMart. The canonical and full set of SS datasets were created in the same manner as zebrafish. The training, weight training, and validation sets were selected from the genes not used in the test set in the same manner as human, and the test set was made from all the nonparalogous genes on chromosomes 2L, and 3L, resulting in a set of 1,938 genes.

For silkworm moth, we downloaded splicing annotations and paralog status for Bmori_2016v1.0 from EnsemblMetazoa’s BioMart and defined the canonical transcripts using the Ensembl canonical set. For this genome, we filtered out genes which were annotated as mitochondrial rather than filtering for genes annotated as chromosomal and made the training, SRE weight learning, and validation sets from the genes not used in the test set as described for human. The test set was made from the non-paralogous genes with the same filters as human on the primary assemblies named BHWX01000012.1, BHWX01000013.1, BHWX01000018.1, BHWX01000021.1, BHWX01000022.1, BHWX01000027.1, BHWX01000038.1, BHWX01000074.1, and BHWX01000097.1, resulting in a set of 920 genes.

For *Arabidopsis*, we used EnsemblPlant’s BioMart to download the paralog status and splicing annotations for genome version TAIR10. As with silkworm moth, we used the Ensembl canonical set to define the canonical transcripts. The training, SRE weight learning, and validation sets were selected from the genes not used in the test set as described for human. The test set was made of non-paralogous genes on chromosomes 2 and 4, resulting in a set of 1,117 genes.

### New MaxEntScan models

To create the training and testing data for our third-order model of human SS, we downloaded the GCF_000001405.39 NCBI assembly for hg38/GRCh38. For each intron-containing, protein-coding gene at least 300 bases in length (to ensure sufficient null examples), we used all 5’SS and 3’SS for true examples, and then randomly selected thirty random non-SS positions for each SS to act as null examples. We removed any sequences which contained non-canonical bases, randomly selected one third of each set to act as test data for Figure 1B, then used the remaining two thirds to train on.

To generate new SS models, we reimplemented the iterative scaling procedure described in the original MaxEntScan paper (Yeo & Burge, 2004). For the 5’SS we used the same 9mer around the SS, but rather than considering the consensus dinucleotide and remaining positions separately, we used the entire 9mer sequence. Similarly for the 3’SS we incorporated the consensus dinucleotide into the considered subsequences. So, in a similar notation as the original paper, where the subscripts refer to sequence position, the probability of generating a 23-nt potential 3’SS sequence *X* is:

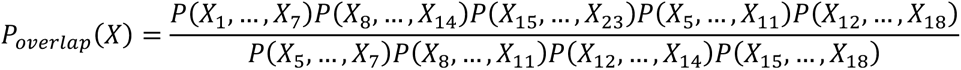

Furthermore, for both the 5’SS sequence and the 3’SS subsequences, the constraints on the maximum entropy distributions include all third-order frequency constraints – i.e. the frequencies of triples of nucleotides at all possible sets of three consecutive or non-consecutive SS positions – in addition to the second and first order constraints. The original MaxEnt methods are used in this paper only in the comparison analyses shown in Figures 1B and 3A.

For the other organisms, the same training procedure and constraints were used, with the training data coming from the all-SS datasets for all genes at least 300 bp in length that are not present in the respective test sets described above. (No validation set is needed for SS training since the parameters are obtained through a deterministic iterative scaling procedure.) These test sets were then used in the comparison analysis in Figure 3A.

### Iterative Learning of Scores

To iteratively learn SRE scores, we began by dividing training set genes (excluding those > 200 kb in length) into four equally sized subsets (plus or minus one gene). For the first subset, we made predictions for the set, either using the CASS model or the SMsplice model. Comparing these predictions to the canonical annotations allowed us to define sets of false positives and false negatives. We counted the occurrences of *k*mers in the intronic and exonic flanking regions for these sets in the same manner as the decoy SS flanking regions described above. This yielded four sets of kmer counts: exonic regions flanking false negatives (*C^e^* ^fn^), exonic regions flanking false positives *C^e^* ^fp^), intronic regions flanking false negatives *C^i^* ^fn^), and intronic regions flanking false positives *C^i^* ^fp^). We then added pseudocounts as follows:

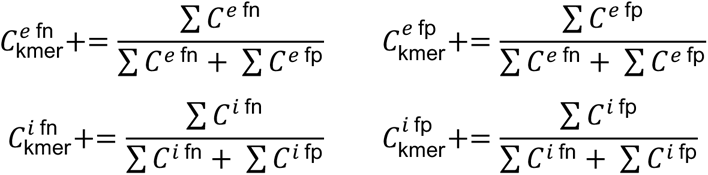

Pseudocounts were added to smooth the empirical frequencies and to ensure that when we normalize each class by the total count to obtain the frequency, any hexamer that did not appear in either the exonic regions flanking false negatives or the exonic regions flanking false positives will be assigned the same frequency for both sets, and likewise in the intronic case. Thus, the log_2_ of the frequency ratio will be zero. This is a desirable property because when we use these resulting frequency ratios to update the SRE scores, we don’t want the scores of kmers that did not appear near false predictions to be affected. Letting the frequencies for a particular kmer be represented By 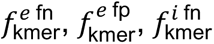, and 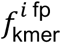; the update is done as follows:

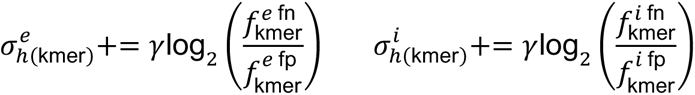

Here *γ* is a learning rate. We used to *γ* = 0.01 throughout this study, because this value performed best in early tests.

### Length Distributions

We used two general methods of smoothing empirical length distributions to create the length distributions for SMsplice. The first method was Gaussian kernel smoothing, accomplished using the neighbors.KernelDensity class of scikitlearn. The second method was adaptive width KDE smoothing, for which we used the GaussianKDE class in the awkde package available at https://github.com/mennthor/awkde with an alpha value of 1 and otherwise default parameters. For both of these methods, we modified the tail of the smoothed distribution in order to allow introns and exons of arbitrarily long lengths. This was done by taking the density of the distribution past a cutoff point, and replacing the tail with an appropriately scaled geometric distribution. So if the value of the distribution at the cutoff point is *p*, the value of the distribution for the next integer length is *p* ⋅ *s*, then for the next integer length it is *p* ⋅ *s*^2^ and so on, where *s* is chosen so that the total density still sums to one. For the intronic distributions, we additionally imposed a minimum length, forcing the probabilities of all the lengths below a certain value to be zero, after which we renormalized the distribution. As all the genes considered in our analyses contained introns, we did not learn length distributions for single-exon genes. The choices of smoothing parameters below were made in each case to improve the fit to the empirical distributions on the training sets, as judged by eye.

For human, the intronic length distribution was smoothed via Gaussian kernel smoothing with a bandwidth 15, a steric constraint of 60, and transitioning to a geometric tail at a length of 1000 nt. All the exonic length distributions were smoothed using adaptive width KDE smoothing, and transitioning to a geometric tail at the 80th percentile of the empirical lengths.

For mouse, exon and intron length distributions were smoothed as in human.

For zebrafish, the intronic length distribution was smoothed via Gaussian kernel smoothing with a bandwidth 5, a steric cutoff of 50, and a geometric tail at 5000. All the exonic length distributions were smoothed using adaptive width KDE smoothing, and a geometric transition at the 80th percentile of the empirical lengths.

For fly, all the length distribution were smoothed via Gaussian kernel smoothing. The intronic length distribution used a bandwidth 5, a geometric tail transition of 2000, and a steric cutoff of 40. The first exon length distribution used a bandwidth 30 and a geometric tail transition of 300. The internal exon length distribution used a bandwidth 30 and a geometric tail transition of 500. The last exon length distribution used a bandwidth 100 and a geometric tail transition of 750.

For moth, the intronic length distribution was smoothed via Gaussian kernel smoothing with a bandwidth 15, a geometric tail transition of 3000, and a steric cutoff of 60. The first and internal exonic length distributions were smoothed using adaptive width KDE smoothing and a geometric transition determined at the 80th percentile of the empirical lengths. The last exon length distribution was smoothed using adaptive width KDE smoothing and a geometric transition determined at the 85th percentile of the empirical lengths.

For *Arabidopsis*, the intronic length distribution was smoothed via Gaussian kernel smoothing with a bandwidth 5, a geometric tail transition of 200, and a steric cutoff of 60. All the exonic length distributions were smoothed using adaptive width KDE smoothing, and a geometric transition determined at the 80th percentile of the empirical lengths.

### SMsplice and Relation to HSMM models

The SMsplice model is most easily described in relation to an HSMM associated with the hidden structure shown in Figure 2A. This structure involves seven hidden states: *E_S_*, *E_F_*, *E*., *E_M_*, *I_L_*, 5′, and 3′, with *S* and *ɛ* representing the traditional start and end states, from which all valid parses must begin and end, respectively. The transition probabilities between states are all set to 0 or 1, as indicated by the arrows, except the transitions from 3′ to *E*_*L*_ and *EO*., which are set to *p_E0_* and *p_E1_* = 1 – *p_E0_*, respectively, where *p_E0_* is the parameter fit in Figure 2B. Here, we have set the *S* to *E_F_* transition probability, *p_ME_*, to 1 and *p_1E_* to 0, effectively assuming that genes have at least one intron, but these transitions can be set differently if one wishes to include intronless genes as a possibility. The length distributions for each of the hidden states were learned as described above; the 5’ and 3’ SS states were assigned a length of 1 with probability 1 for convenience, though of course the associated core SS motifs extend upstream and downstream of these positions.

To fully specify a standard HSMM would require a set of possible emissions and probability distributions over these emissions for each hidden state, and one could then find the highest probability parse for any sequence of emissions using the HSMM version of the Viterbi algorithm (Shun-Zheng & Kobayashi, 2003). However, our main goal here was to provide a framework to apply the structural constraints of the spliceosome, particularly length distributions and SS pairing, for use in discriminating more and less plausible splicing patterns, rather than building a generative model. Therefore, where an HSMM would consider the probability of some parse *π* for a given sequence, ℙ[seq, *π*], for SMsplice we define *SM*[seq, *π*] an expression analogous to the base-2 logarithm of ℙ[seq, *π*], normalized to a background model of sequence composition. For example, our SS-only model uses the MaxEnt log-odds ratios (representing the log of the probability of generating a sequence segment under the SS model to that of generating it under a background model) in place of the terms an HSMM would use for emissions from the SS hidden states. And it assigns a log-odds value of 0 in place of the emissions for other states (representing introns and different types of exons), effectively treating these states as not different from background (i.e. odds-ratio of 1). The CASS model makes similar substitutions, replacing SS emission terms by CASS scores, which derive from log-odds ratios but do not necessarily correspond to the log-odds ratios of any specific pair of sequence-generative models; again, the emissions terms for other states are replaced by log-odd values of 0, effectively ignoring the composition of exons and introns outside of the local regions that contribute to CASS scores.

Because the SMsplice model is defined in log-odds rather than generative terms, it is technically a semi-Markov CRF (Lafferty et al., 2001; Sarawagi & Cohen, 2004). It satisfies the Markov condition that the odds of hidden state *i*+1 given seq and any subset of hidden state values depends only on seq and the previous state *i* (via the transition probability from the state *i* to state *i*+1). The CASS framework defines the SS score at some position *j* as a function of sequences up to ∼100 bp upstream and downstream of j (via the core SS motif, ESR and ISR scores). Such a definition is compatible with a CRF framework, where conditional probabilities (or log-odds) of hidden states are defined conditionally on the entire input sequence.

Consider a parse *π* of a sequence of length *T* which has *N* > 0 introns of lengths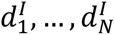, and therefore *N* − 1 internal exons which have lengths 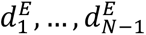. Let the first exon have length 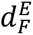 and the last exon have length 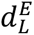. Then, otherwise using the notation discussed above and shown in Figure 2, the base 2 logarithm of the complete data likelihood of the fully specified HSMM would be:

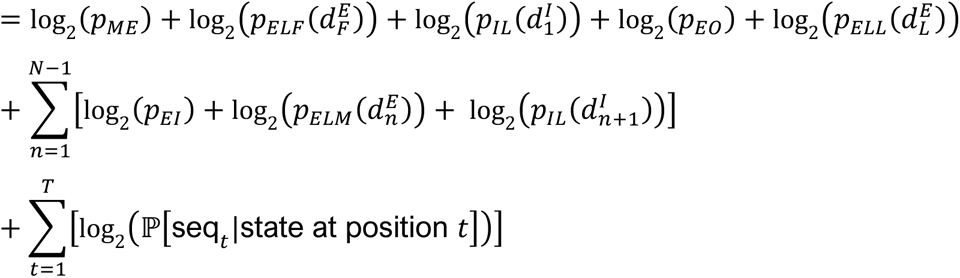

The SMsplice function is reproduced here, with minor modifications to emphasize correspondence with the HSMM:

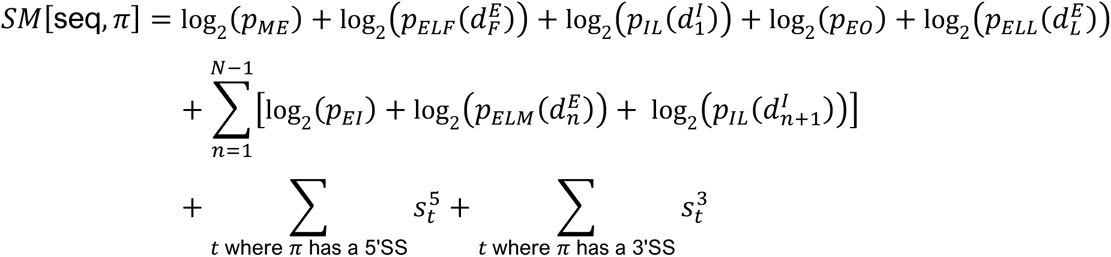

Thus, *SM*[seq, *π*] uses the same sums of log-probabilities as in an HSMM, except that the emissions term is replaced by CASS scores, which involve log-odds of SS-associated sequences relative to background or decoy sequences. Therefore, SMsplice is a discriminative model whose maximum can be found by applying the Viterbi algorithm for the associated HSMM. The parse *π*^∗^ which maximizes *SM*[seq, *π*] is taken to be the SMsplice prediction for the input sequence.

### Decoy SS set and Real Versus Decoy Scores

To identify decoy SS, we scored every position in transcripts of the training set using the third-order MaxEntScan methods for the organism in question. For 5’SS and 3’SS, we collected the scores of all the positions annotated in the canonical splicing pattern which flanked an internal exon. For each of these real SS scores, ordered from strongest to weakest, we selected a random gene in the training set. If that gene had a position that was not annotated as a splice site (from gtex_dataset.txt for human and from the all-SS datasets for other organisms), was not already selected as a decoy, and had an associated MaxEnt score within 0.5 bits of the real SS in question, we designated that position as a decoy. If the gene lacked a site with appropriate score, another gene was selected at random without replacement. If none of the genes in the training set had such a position, the SS was left unmatched. All SS were matched by decoys for human, mouse, zebrafish and moth, while fly had just 13 unmatched 3’SS (less than 1%) and 1,351 unmatched 5’SS (11%), and *Arabidopsis* had 2,131 unmatched 3’SS (11%) and 1,610 unmatched 5’SS (8%).

To score *k*mers using the matched decoy SS set, we counted the occurrence of each *k*mer in the flanking regions, the regions of length *r* which abut but do not overlap the MaxEntScan regions. Flanking regions were also required to remain within the boundaries of the gene; if after this shortening the region was reduced to less than *k* bases, the region was discarded. The *k*mer counts for real SS were similarly calculated, further shortening the flanking region to avoid overlap with any other SS motifs in the vicinity, if necessary (considering the segment scored by MaxEntScan to represent the SS motif). So, for instance, in the case of an exon of length < *r*, the flanking region upstream of the 5’SS and downstream of the 3’SS would be the same exonic sequence. Overall, this procedure yielded *k*mer counts upstream and downstream of real and decoy 5’SS and 3’SS. *k*mers upstream of real and decoy 5’SS were considered exonic, while downstream *k*mers were considered intronic, with the opposite convention used for real and decy 3’SS. A pseudocount of two was added to each of these counts for smoothness, and then each category was normalized to obtain the frequency of each kmer in each region.

Representing the resulting frequency values for a particular *k*mer as 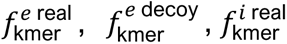 and 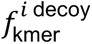, we seeded exonic and intronic scores by assigning *σ^i^* and

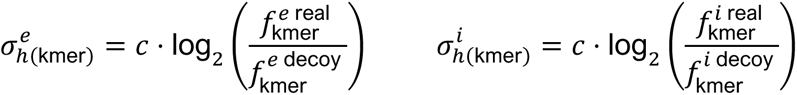

Here, the “SRE weight constant” *c* > 0 was chosen using a binary search to maximize the F_1_ performance on the SRE weight learning set when using these SRE scores. This binary search was initialized by considering the values *c* = 0 and *c* = 1, and our binary search was limited to 16 values. The resulting scores are used in all the relevant analyses following Figure 4B and Supplemental Figure 5.

### Clustering Hexamers

For the clustering analyses shown in Figure 5, we defined the “distance” (dissimilarity) between two *k*mers by checking the agreement of the sequences for each possible amount of overlap. For an overlap of one base, the base distance score was *k* − 1; for an overlap of two bases, the distance was *k* − 2, and so on. Then, for every base that did not agree in the overlapping potion, we incremented the distance by 1. Finally, we defined the distance as the minimum base distance score obtained from evaluating all 2*k* − 1 possible overlaps between the two *k*mers. These distances were used to construct a distance matrix for all the *k*mers in the relevant set, and the kmers were clustered based on this matrix using average linkage hierarchical clustering to define ten clusters of at least four *k*mers. As in the RESCUE-ESE paper, the *k*mers in each cluster containing at least four *k*mers were aligned using ClustalW with default parameters (Fairbrother et al., 2002). A PWM for each cluster was calculated from the frequency of each base at each position in the resulting alignment; these frequencies may sum to < 1 at positions where one or more sequences were not aligned.

### PWM Comparisons

We first pruned any position where fewer than one third of the sequences aligned. We then replaced these positions with a uniform distribution (frequency of ¼ for all 4 bases), and also added padded each PWM with uniform positions so that all PWMs extended for the full extent of the alignment. To compare two PWMs, we considered all possible ungapped alignments of the two PWMs, defining the PWM distance for each specific alignment as the sum of the Jensen-Shannon Divergence (JSD) values between corresponding positions; the distance between the two PWMs was then defined as the minimum PWM distance across all alignments of the PWMs. When clustering the PWMs within each category for all organisms, we used this distance for average linkage hierarchical clustering.

### RNAcompete Comparisons

Using the PWM distance measure described above, we compared each of the PWMs determined for our clusters with each of the RNAcompete PWMs from the relevant organism, and considered the smallest value as the best match. When that best matching motif was associated with multiple proteins, all of those proteins are reported. To determine a significant distance, we considered all of the PWMs for all of the clusters and changed their bases using the permutation A à G à T à C à A, e.g., a PWM with consensus sequence ATCCG would be converted to one with consensus GCAAT. Then we scored these permuted PWMs against all the RNAcompete motifs for the relevant organism, plotted the distribution of minimum distances, and took the first percentile of this distribution as our distance cutoff, corresponding to P < 0.01.

### Cluster Splicing Activity

Fine mapping of sQTLs and eQTLS in GTEx was performed by Barbeira and coworkers using the DAP-G fine-mapping method (Barbeira et al., 2019). We accessed this dataset using their zenodo link. Note that fine-mapping was only done on European Ancestry samples of GTEx. We filtered for intron clusters with two or three introns per cluster, in order to filter for splicing events that can be meaningfully described as either alternative splice site exons or skipped exons. We created pan-tissue clusters by merging the locations of each intron in a cluster, dropping tissue-specific cluster IDs. We then sorted by posterior inclusion probability (PIP) values in descending order, and dropped duplicates based on pan-tissue cluster IDs and variant pairs, keeping the first value. This insured that we obtained the top variant-phenotype pair for all splicing events.

From this set, we further filtered for only SNP variants and splicing events that represented skipped exons and choices between exactly two alternative 5’SS or 3’SS. For each of these events, we considered only the variants that fell in the regions that would be relevant to the SRE contribution to CASS scores of any of the SS, using the regions it fell in to determine if the variant was intronic, exonic, or both. For each cluster, we determined the set of relevant variants by selecting those that fell in a relevant CASS region of an annotated SS and for which the (reference or alternative) variant fell within a hexamer of the cluster, and changed the score of the resulting hexamer to the opposite sign. For instance, for an ESE cluster we looked for variants that fell in exonic CASS regions where the variant overlapped a component hexamer of that cluster, and the variant changed the hexamer to one with an ESR score < 0.

We determined a control set of variants by considering variants that fell in CASS regions but did not fall in any of the most extreme 5% hexamers, i.e. none of the hexamers which overlapped the variant in either condition were in the top or bottom 5% of ESRs or ISRs. We then compared the distributions of PIP values associated with this control set of variants to those associated with each of the clusters. If the clusters are indeed associated with splicing regulation, we would expect their PIP values to be generally larger than those for the control set. To measure this, we used a one-sided Kolmogorov-Smirnov (KS) test to assess whether the distributions were significantly shifted in the appropriate direction, indicating increased likelihood of causality. We used Benjamini-Hochberg multiple test correction with alpha = 0.1 to determine final significance shown in Figure 5C.

## Supplemental Figures

**Supplemental Figure 1:**
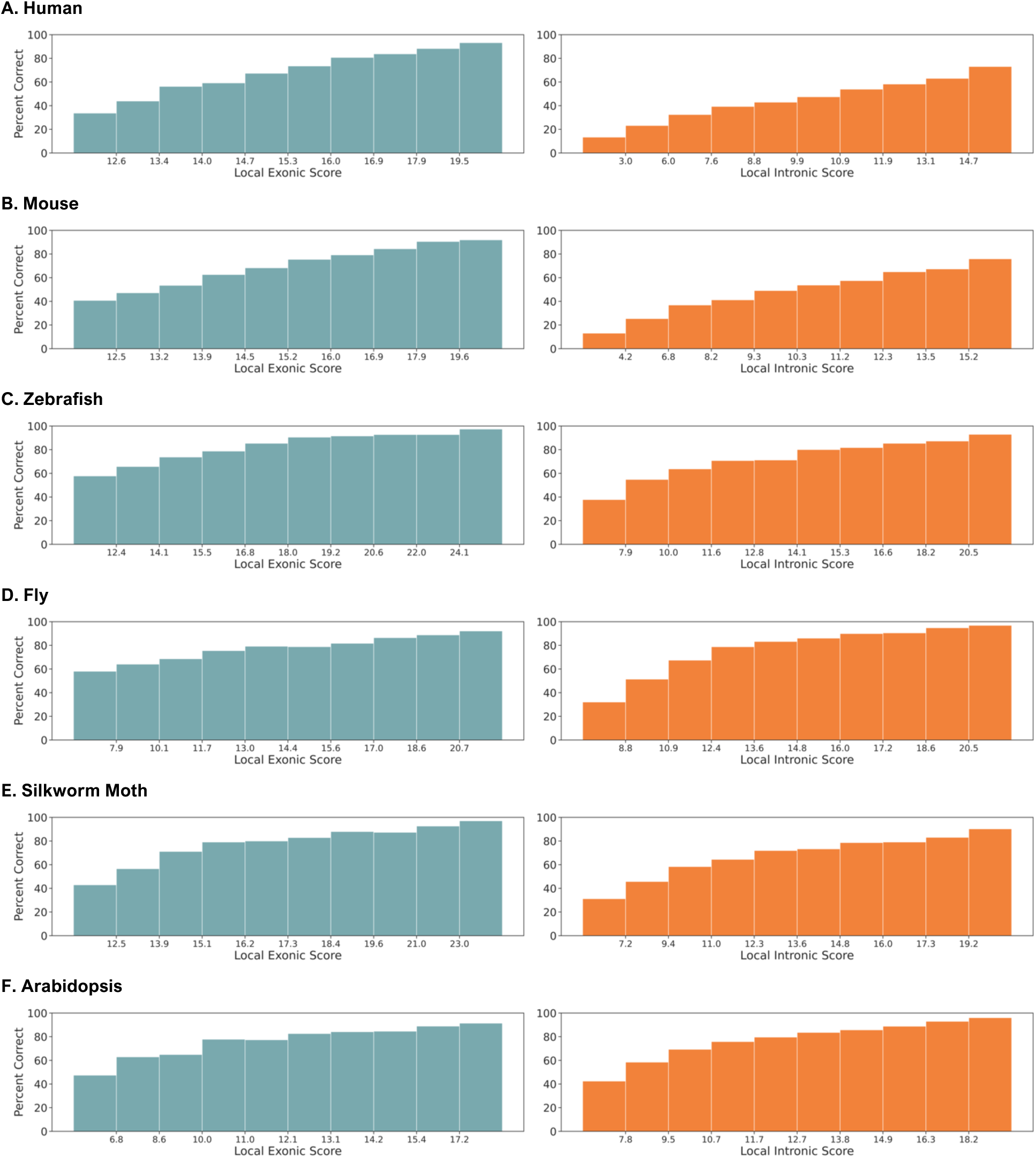
Accuracy increases with local ccore for exons and introns. Percent correct for predicted internal exons and introns split into 10 quantile bins of local exonic and intronic scores for A) human, B) mouse, C) zebrafish, D) fly, E) moth, and F) *Arabidopsis*.

**Supplemental Figure 2:**
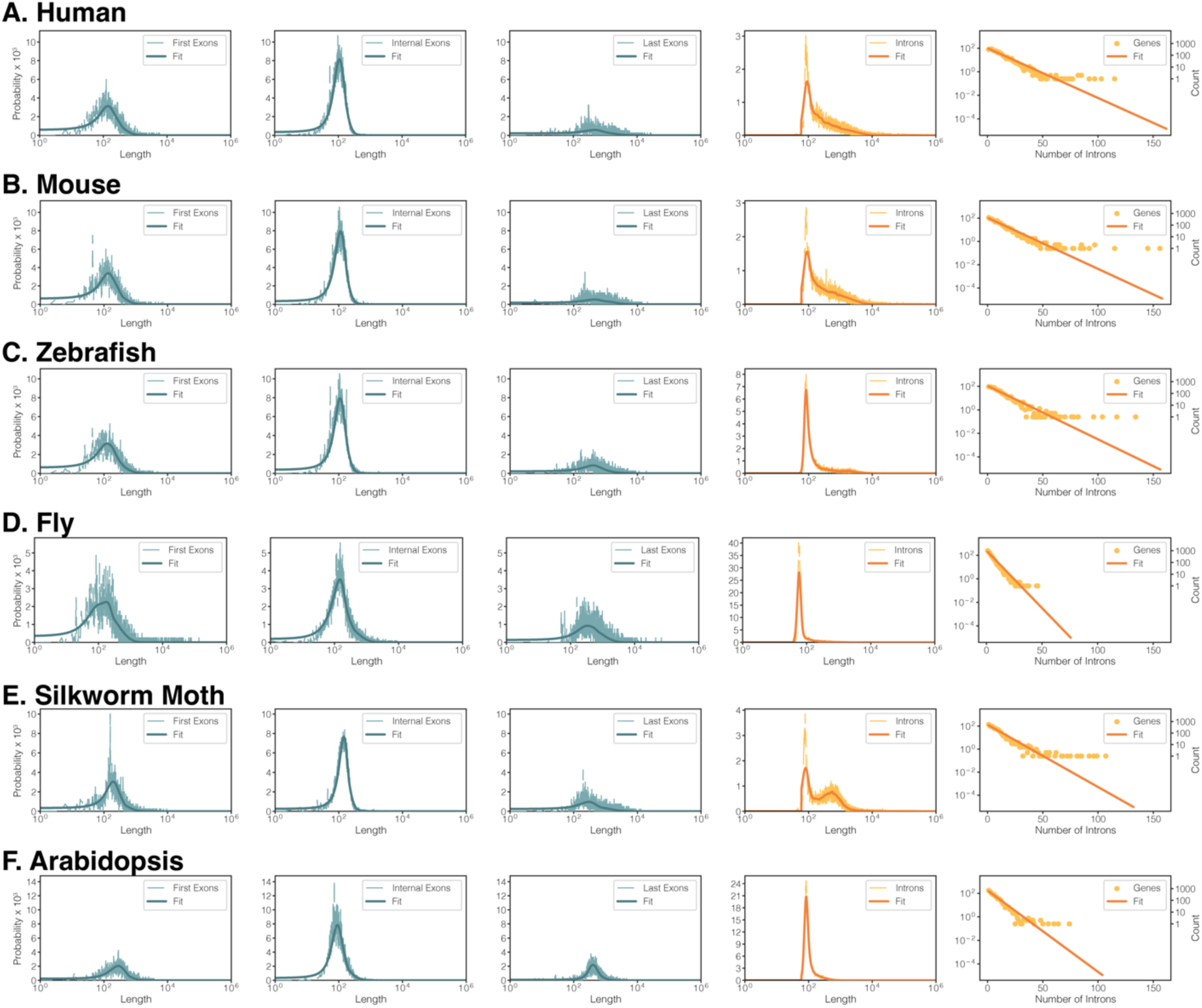
Exon and Intron length distributions and intron number per gene. Length distributions and number of introns distribution used in the SMsplice model for A) human, B) mouse, C) zebrafish, D) fly, E) moth, and F) *Arabidopsis*, along with the empirical distributions they were fit to.

**Supplemental Figure 3:**
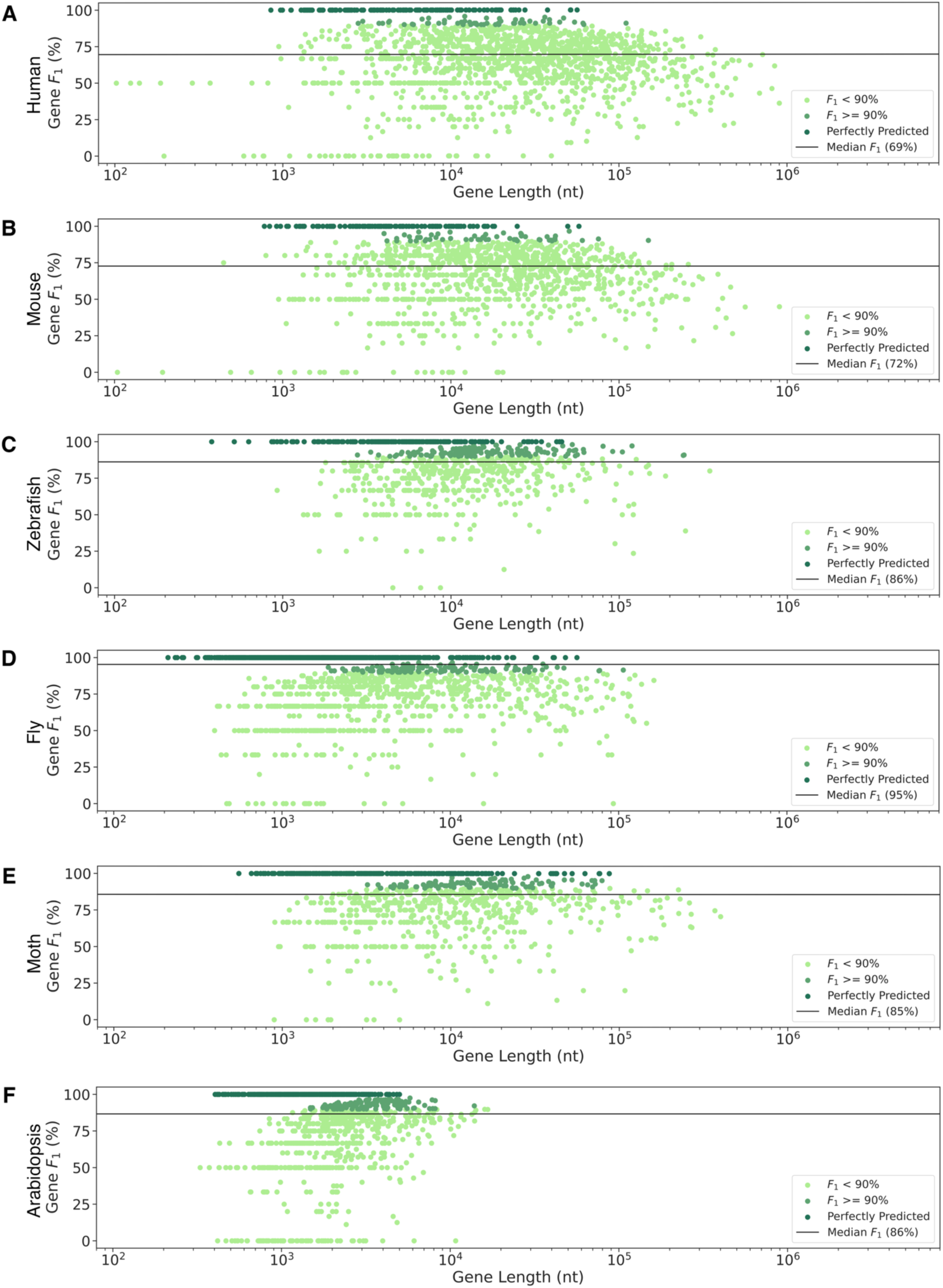
Individual SMsplice Gene Predictions. Gene-level F_1_ accuracy for the test sets of A) human, B) mouse, C) zebrafish, D) fly, E) moth, and F) *Arabidopsis* genes. Each dot represents the length (x-axis) and accuracy (y-axis) for a single gene, with the median gene-level F_1_ accuracy shown by a horizontal line. Dots are color-coded by accuracy bin (< 90%, light green; > 90%, medium green; perfectly predicted, dark green) to facilitate visualization.

**Supplemental Figure 4:**
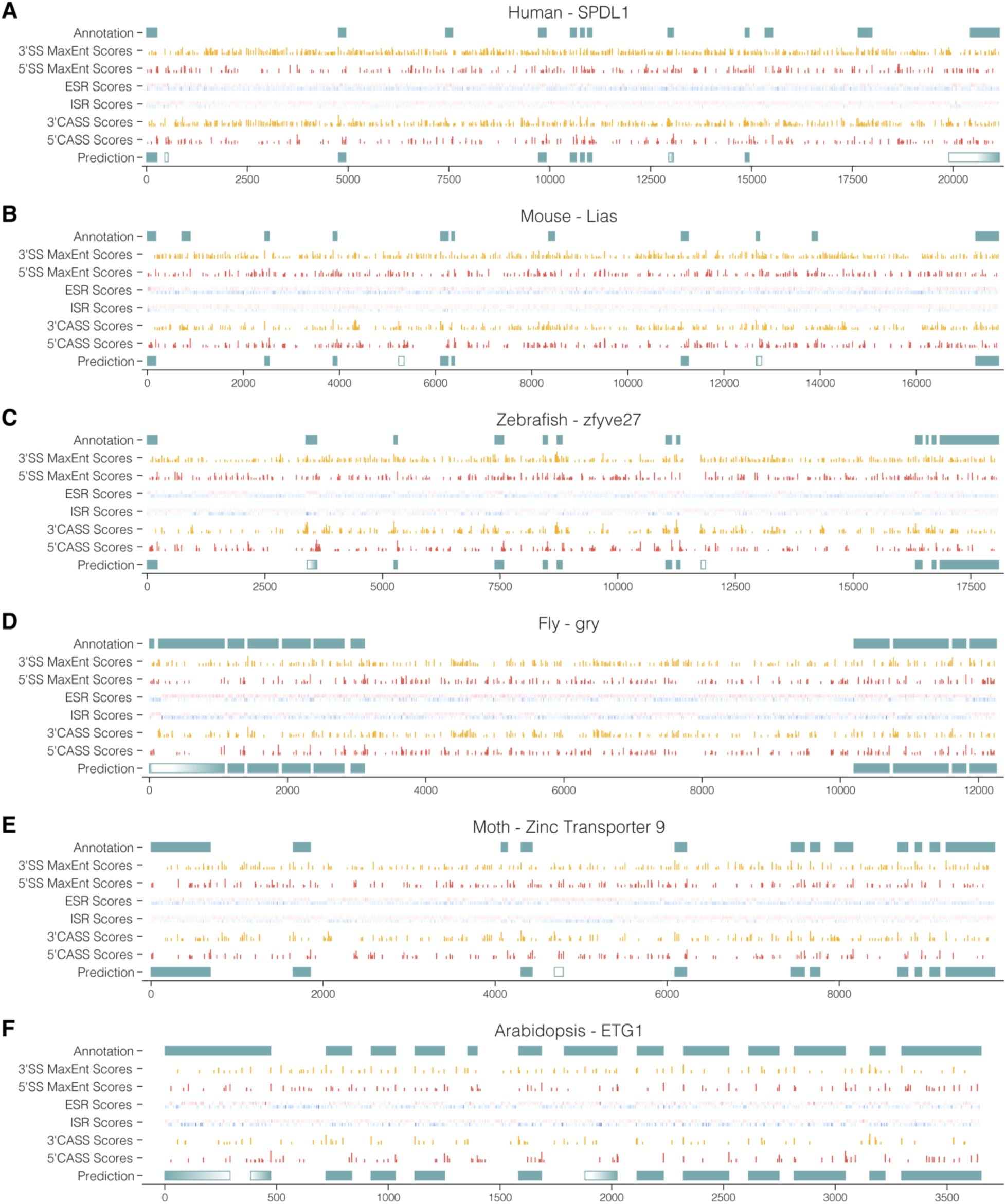
Example Predictions for each organism. An illustration of the SMsplice predictions for each organism, with the gene chosen to have a prediction with an F_1_ score within a percentage of the median for that organism. A) Human gene SPDL1 with F_1_ 70%. Mouse gene Lias with F_1_ 72%. C) Zebrafish gene zfyve27 with F_1_ 86%. D) Fly gene gry with F_1_ 95%. E) Moth gene Zinc Transporter 9 with F_1_ 86%. F) *Arabidopsis* gene ETG1 with F_1_ 88%.

**Supplemental Figure 5:**
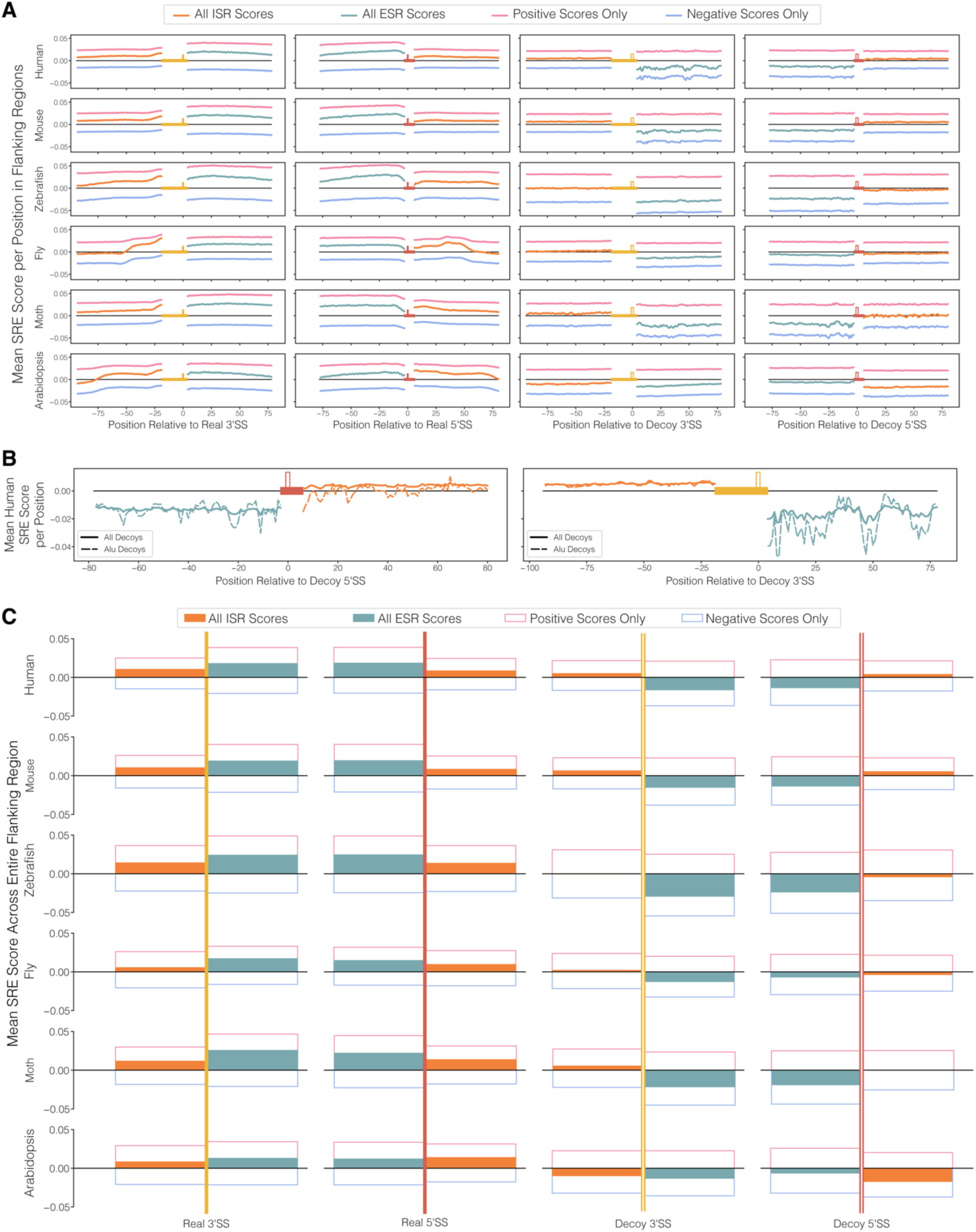
Distribution of SREs in flanking regions of real and decoy SS. A) The relevant SMsplice-learned ESR and ISR scores, along with their positive and negative components, for each position in the flanking regions, averaged separately across all real and decoy 5’SS and 3’SS in the training set for each organism. B) The relevant SMsplice-learned ESR and ISR scores for each position in the flanking regions, averaged separately across all decoy 5’SS and 3’SS in the training set as well as for the subset of these decoy SS that intersect Alu elements. C) The average of the scores shown in A) across the associated regions for each of the organisms.

**Supplemental Figure 6:**
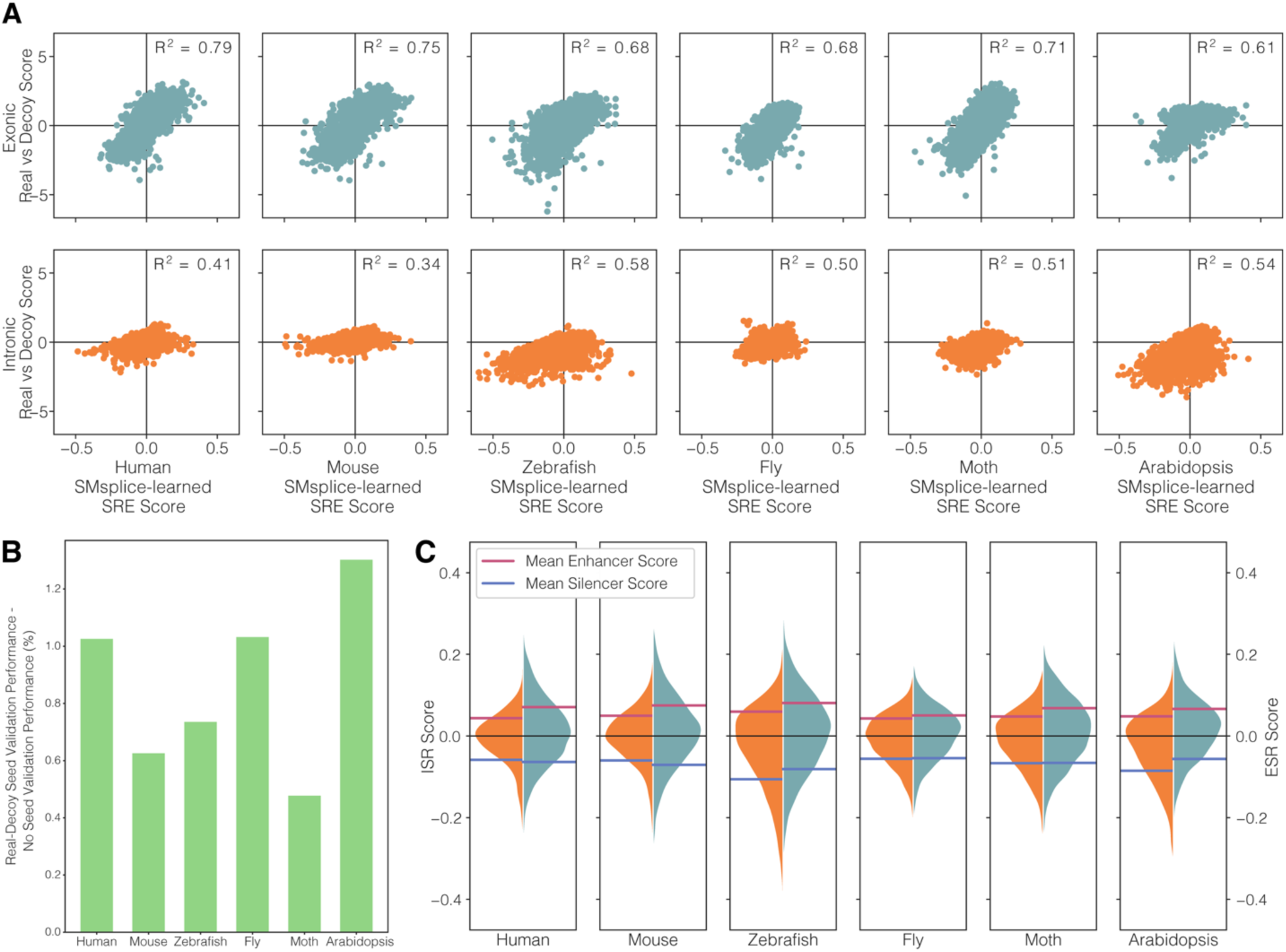
Use of real-vs-decoy SS to seed SRE scores. A) Real versus decoy scores in each organism are positively correlated with their respective SMsplice-learned SRE scores with no seed. B) SMsplice-learned SRE scores with weighted real versus decoys scores acting as a seed outperform the SMsplice-learned SRE scores with no seed for all organisms. C) Density of the SMsplice-learned ESR and ISR scores for each organism as well as the averages of all the positive scores and all the negative scores.

**Supplemental Figure 7:**
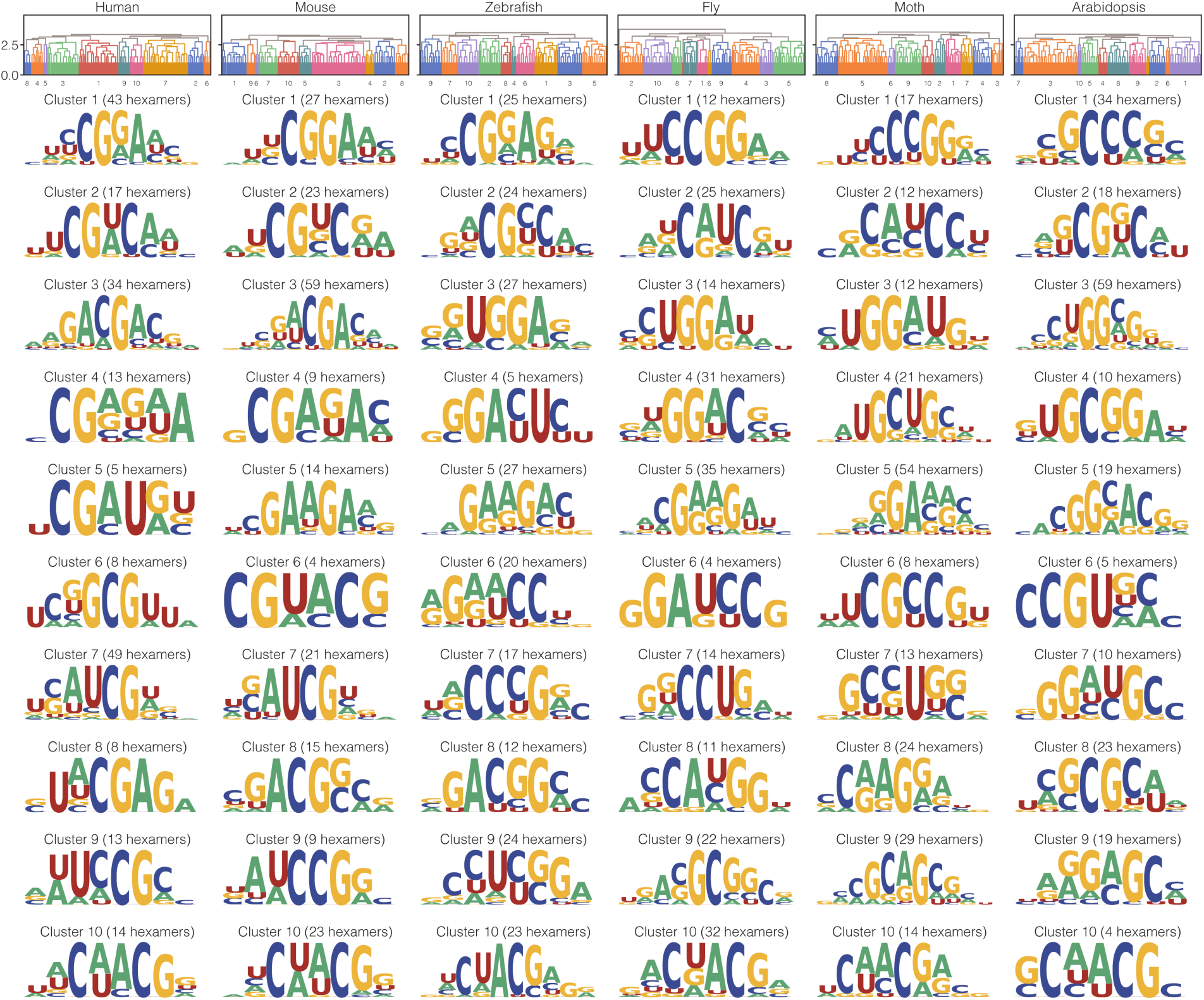
ESE clusters for all organisms. For all the organisms, dendrograms showing the clusters made for the hexamers with the top 5% ESR scores and the motifs created by aligning these clusters, as in Figure 5.

**Supplemental Figure 8:**
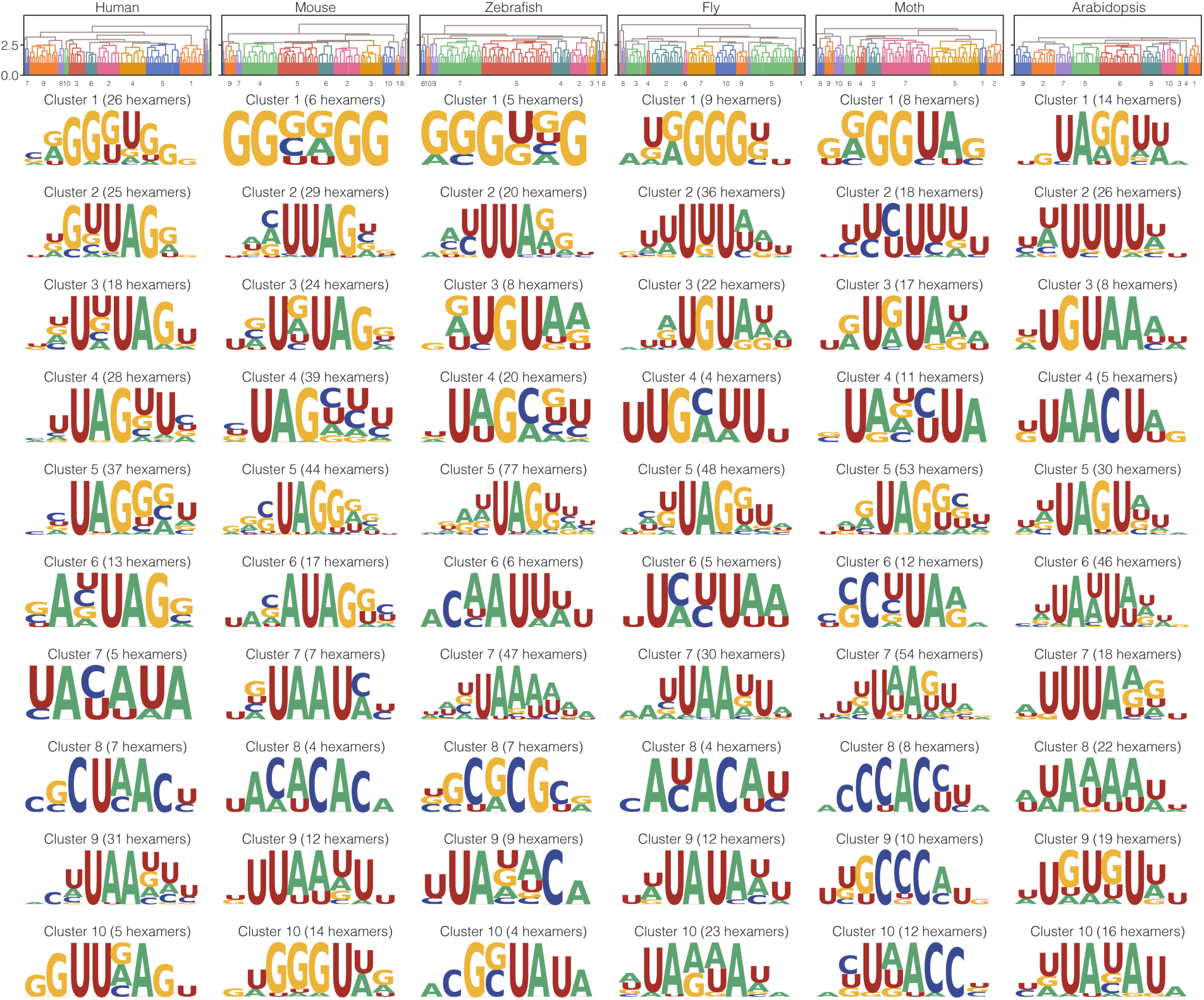
ESS clusters for all organisms. For all the organisms, dendrograms showing the clusters made for the hexamers with the bottom 5% ESR scores and the motifs created by aligning these clusters, as in Figure 5.

**Supplemental Figure 9:**
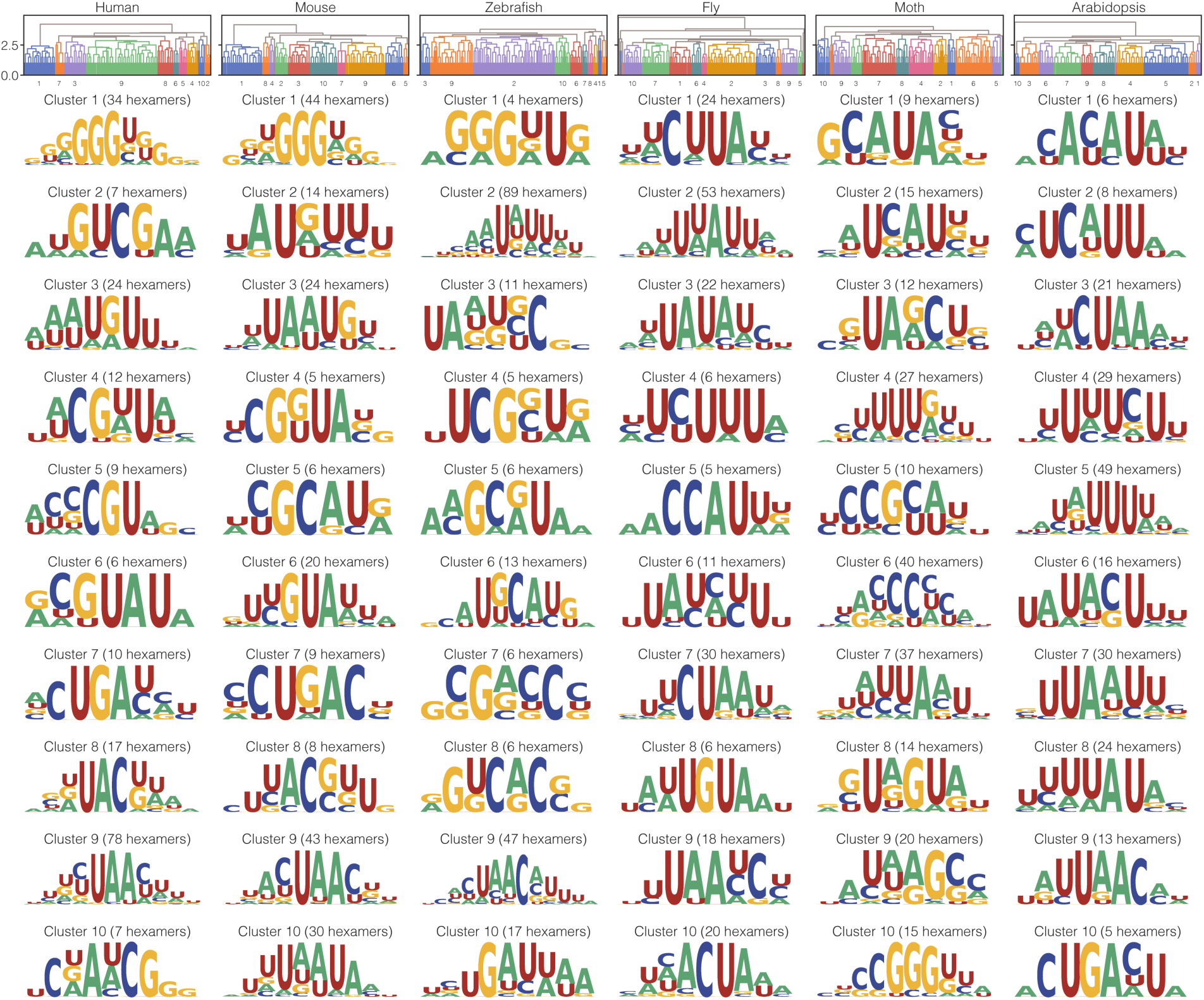
ISE clusters for all organisms. For all the organisms, dendrograms showing the clusters made for the hexamers with the top 5% ISR scores and the motifs created by aligning these clusters, as in Figure 5.

**Supplemental Figure 10:**
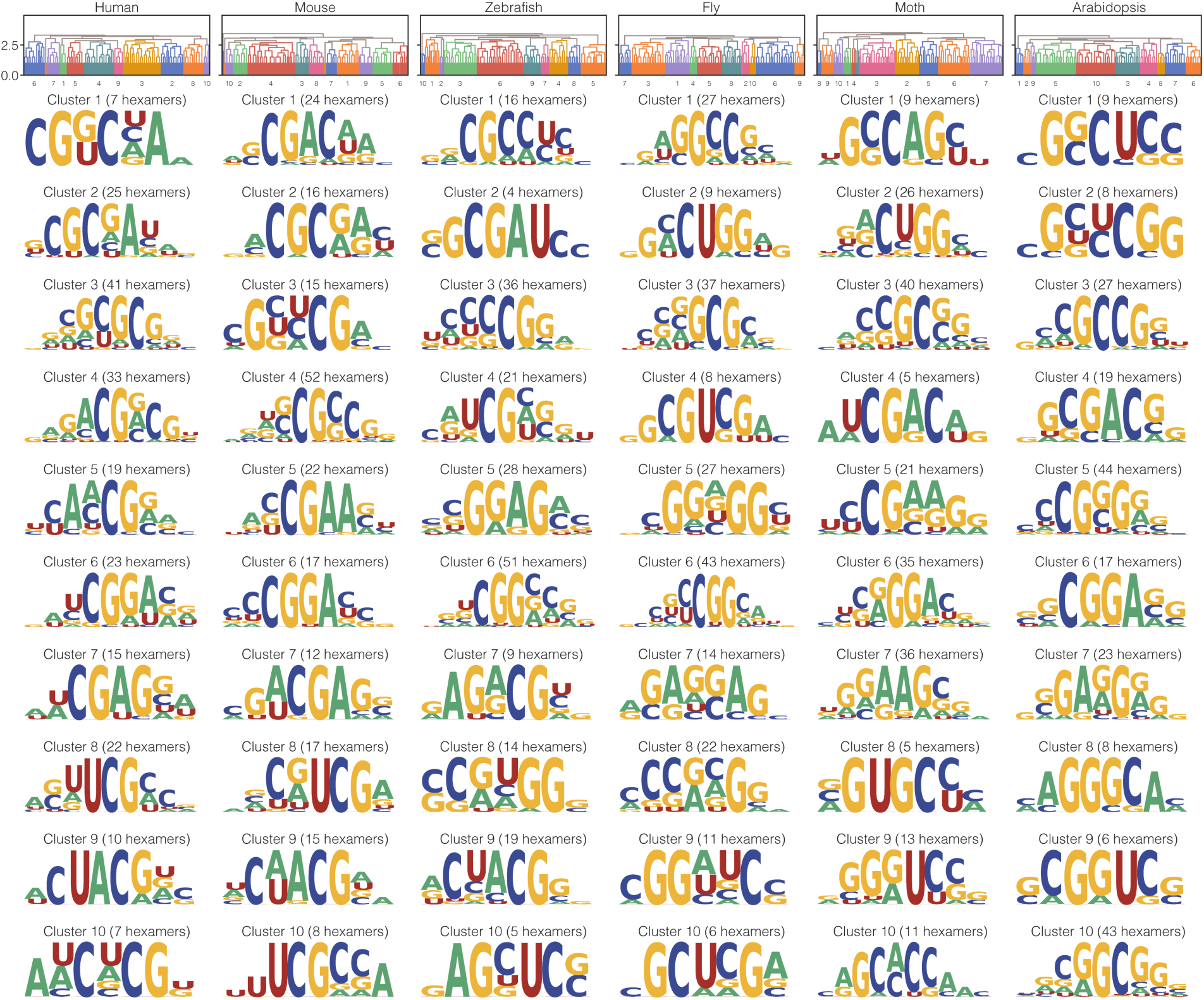
ISS clusters for all organisms. For all the organisms, dendrograms showing the clusters made for the hexamers with the bottom 5% ISR scores and the motifs created by aligning these clusters, as in Figure 5.

**Supplemental Figure 11:**
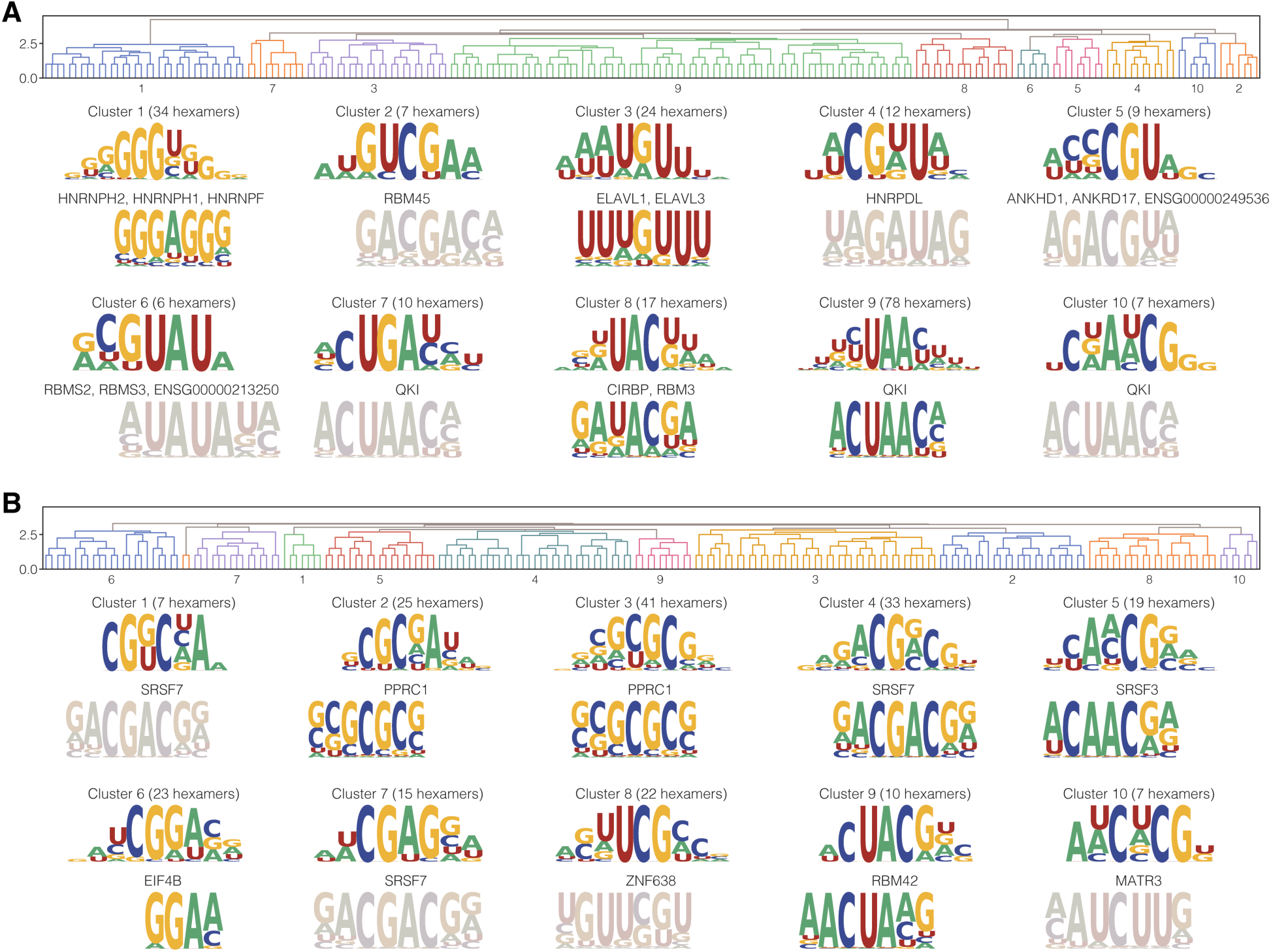
Comparison of human ISE and ISS motifs to human RNACompete motifs. A) Dendrogram for clustering hexamers with the 5% most positive ISR scores in human, as well as the resulting aligned clusters and best RNAcompete matches below those clusters. Desaturation (graying out) indicates that the best match did not satisfy our threshold. B) Dendrogram for clustering hexamers with the 5% most negative ISR scores in human, as well as the resulting aligned clusters and best RNAcompete matches below those clusters.

**Supplemental Figure 12:**
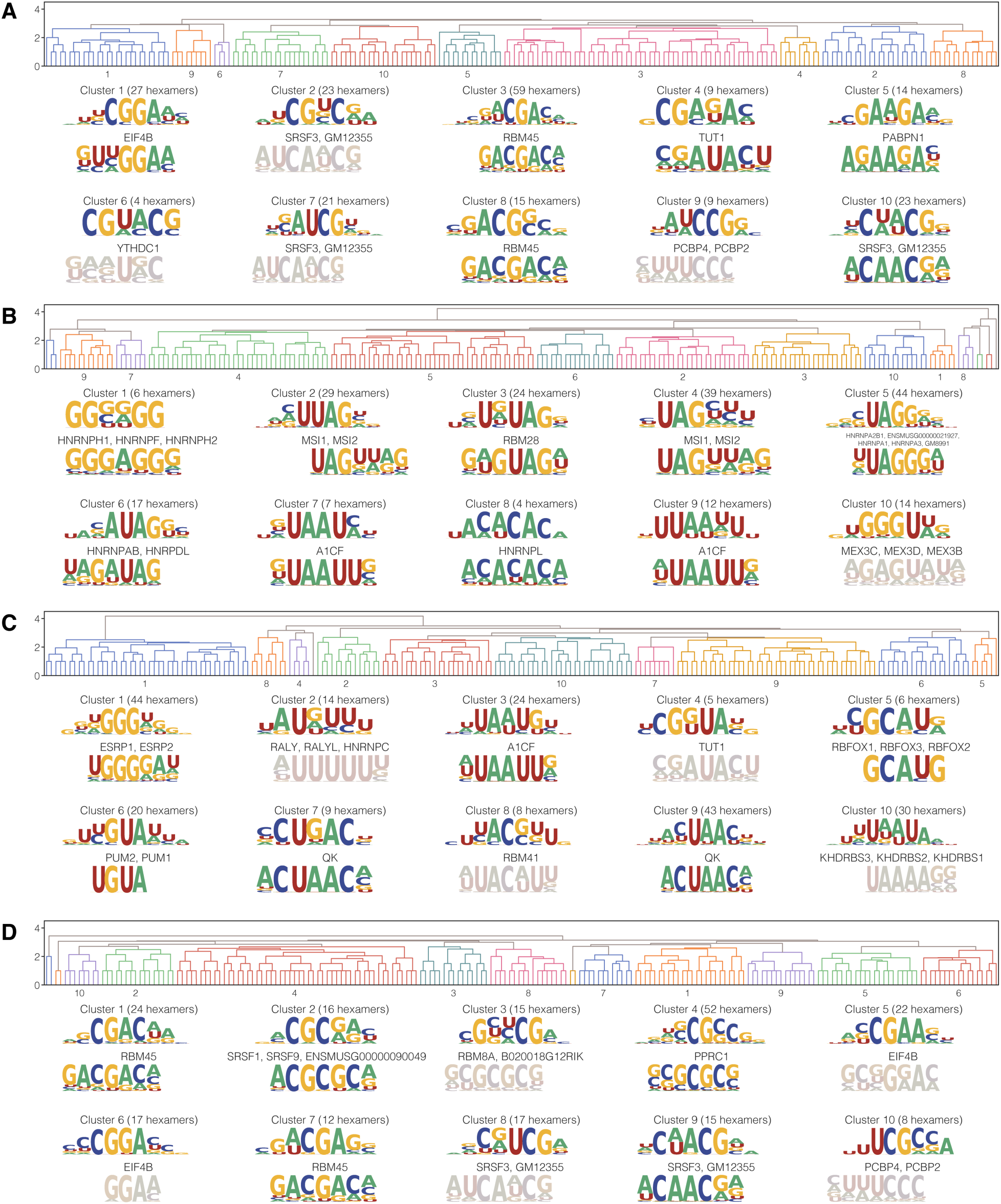
Comparison of mouse SRE motifs to mouse RNACompete motifs. A) Dendrogram for clustering hexamers with the 5% most positive ESR scores in mouse, as well as the resulting aligned clusters and best RNAcompete matches below those clusters. Desaturation (graying out) indicates that the best match did not satisfy our threshold. B) Dendrogram for clustering hexamers with the 5% most negative ESR scores in mouse, as well as the resulting aligned clusters and best RNAcompete matches below those clusters. C) Dendrogram for clustering hexamers with the 5% most positive ISR scores in mouse, as well as the resulting aligned clusters and best RNAcompete matches below those clusters. D) Dendrogram for clustering hexamers with the 5% most negative ISR scores in mouse, as well as the resulting aligned clusters and best RNAcompete matches below those clusters.

**Supplemental Figure 13:**
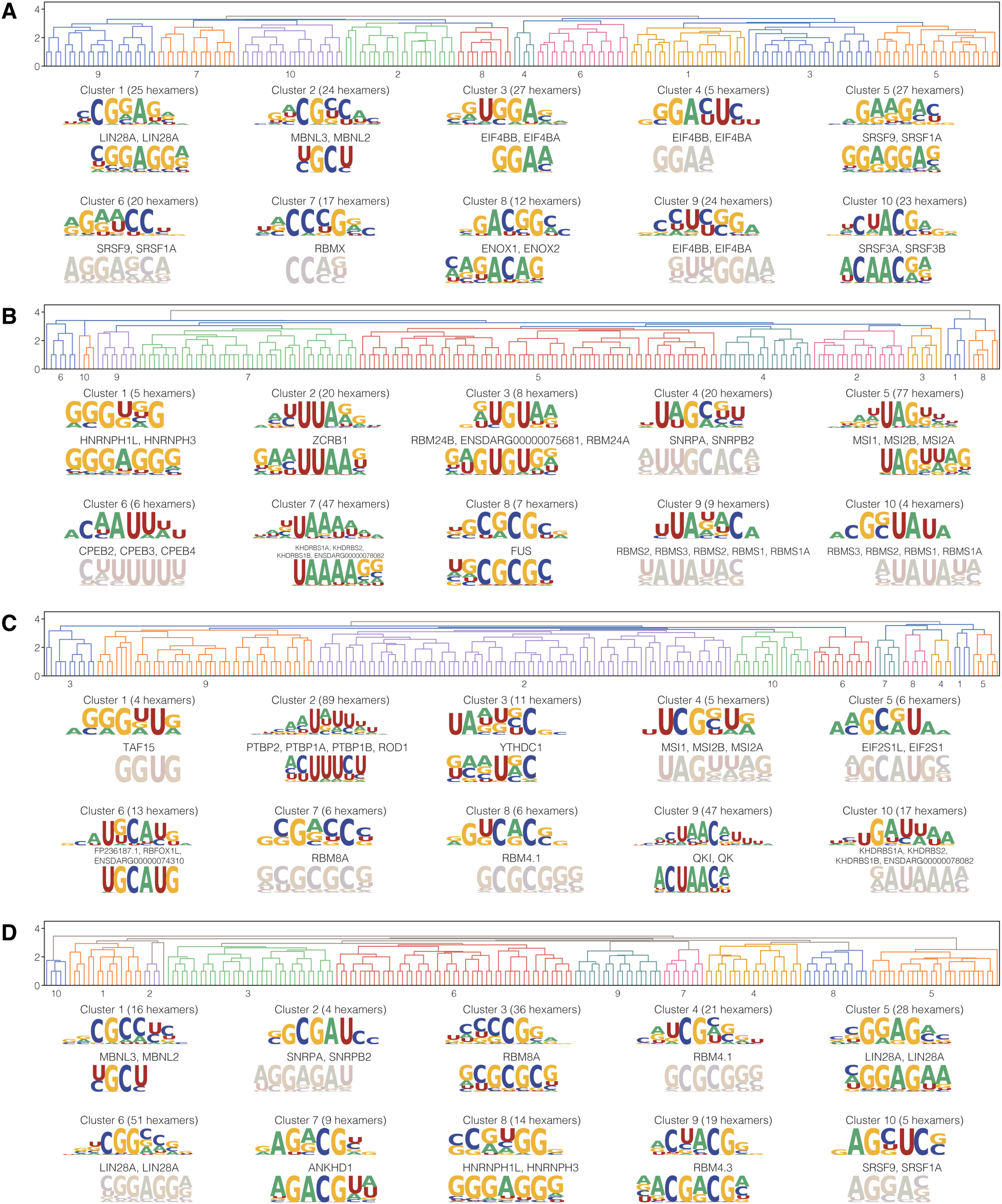
Comparison of zebrafish SRE motifs to zebrafish RNACompete motifs. A) Dendrogram for clustering hexamers with the 5% most positive ESR scores in zebrafish, as well as the resulting aligned clusters and best RNAcompete matches below those clusters. Desaturation indicates that the best match did not satisfy our threshold. B) Dendrogram for clustering hexamers with the 5% most negative ESR scores in zebrafish, as well as the resulting aligned clusters and best RNAcompete matches below those clusters. C) Dendrogram for clustering hexamers with the 5% most positive ISR scores in zebrafish, as well as the resulting aligned clusters and best RNAcompete matches below those clusters. D) Dendrogram for clustering hexamers with the 5% most negative ISR scores in zebrafish, as well as the resulting aligned clusters and best RNAcompete matches below those clusters.

**Supplemental Figure 14:**
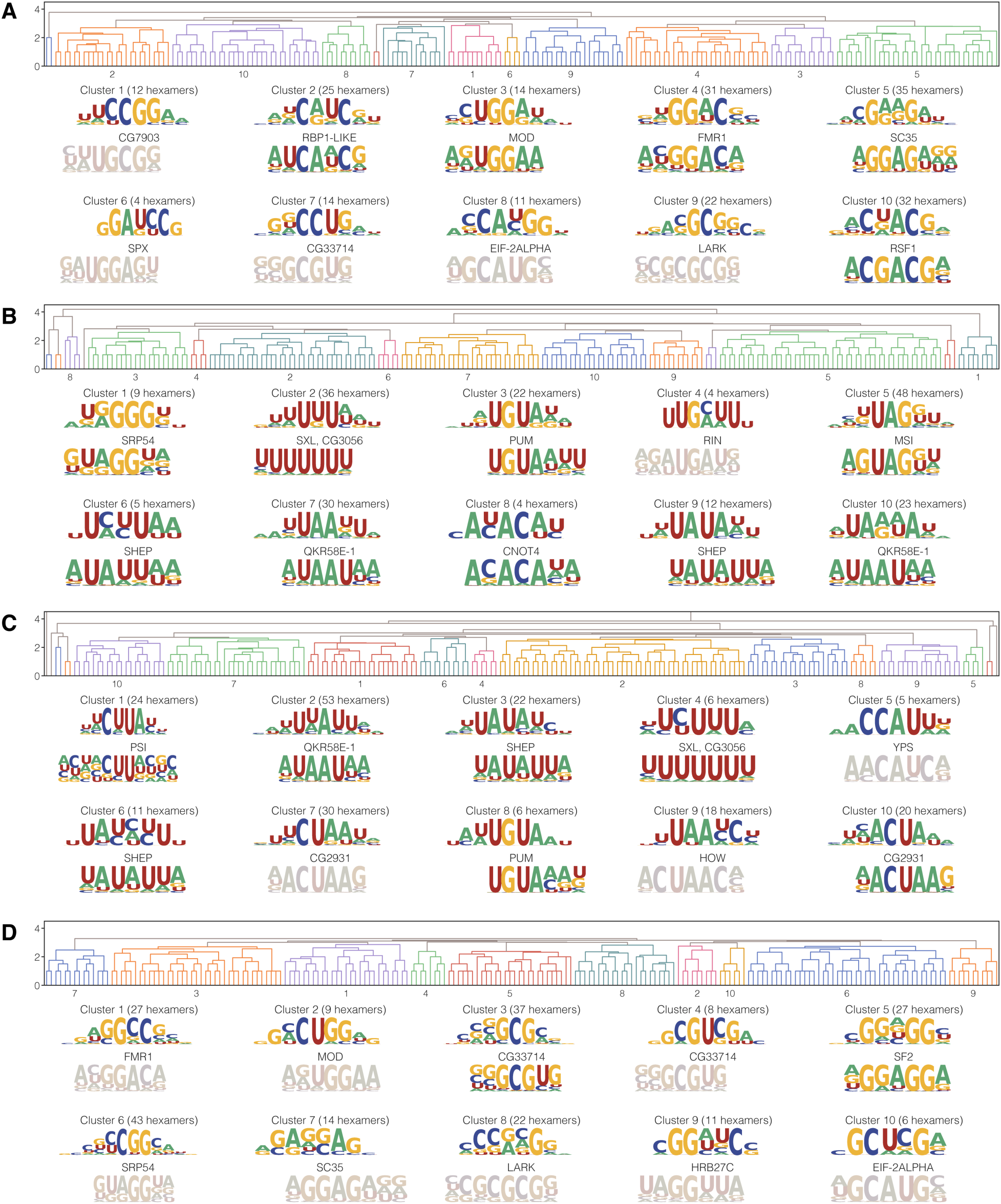
Comparison of Drosophila ESR motifs to Drosophila RNACompete motifs. A) Dendrogram for clustering hexamers with the 5% most positive ESR scores in fly, as well as the resulting aligned clusters and best RNAcompete matches below those clusters. Desaturation indicates that the best match did not satisfy our threshold. B) Dendrogram for clustering hexamers with the 5% most negative ESR scores in fly, as well as the resulting aligned clusters and best RNAcompete matches below those clusters. C) Dendrogram for clustering hexamers with the 5% most positive ISR scores in fly, as well as the resulting aligned clusters and best RNAcompete matches below those clusters. D) Dendrogram for clustering hexamers with the 5% most negative ISR scores in fly, as well as the resulting aligned clusters and best RNAcompete matches below those clusters.

**Supplemental Figure 15:**
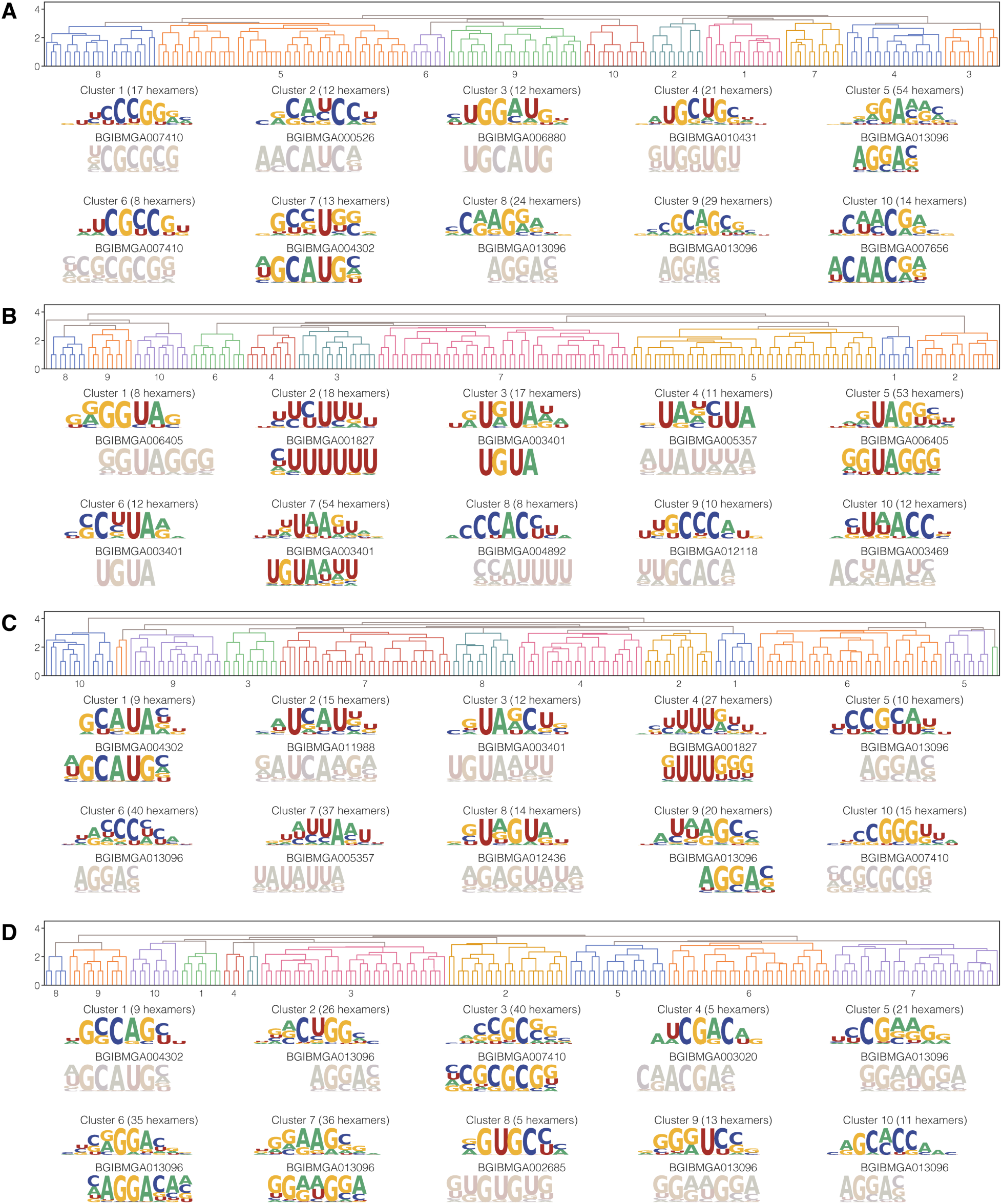
Comparison of moth SRE motifs to moth RNACompete motifs. A) Dendrogram for clustering hexamers with the 5% most positive ESR scores in moth, as well as the resulting aligned clusters and best RNAcompete matches below those clusters. Desaturation indicates that the best match did not satisfy our threshold. B) Dendrogram for clustering hexamers with the 5% most negative ESR scores in moth, as well as the resulting aligned clusters and best RNAcompete matches below those clusters. C) Dendrogram for clustering hexamers with the 5% most positive ISR scores in moth, as well as the resulting aligned clusters and best RNAcompete matches below those clusters. D) Dendrogram for clustering hexamers with the 5% most negative ISR scores in moth, as well as the resulting aligned clusters and best RNAcompete matches below those clusters.

**Supplemental Figure 16:**
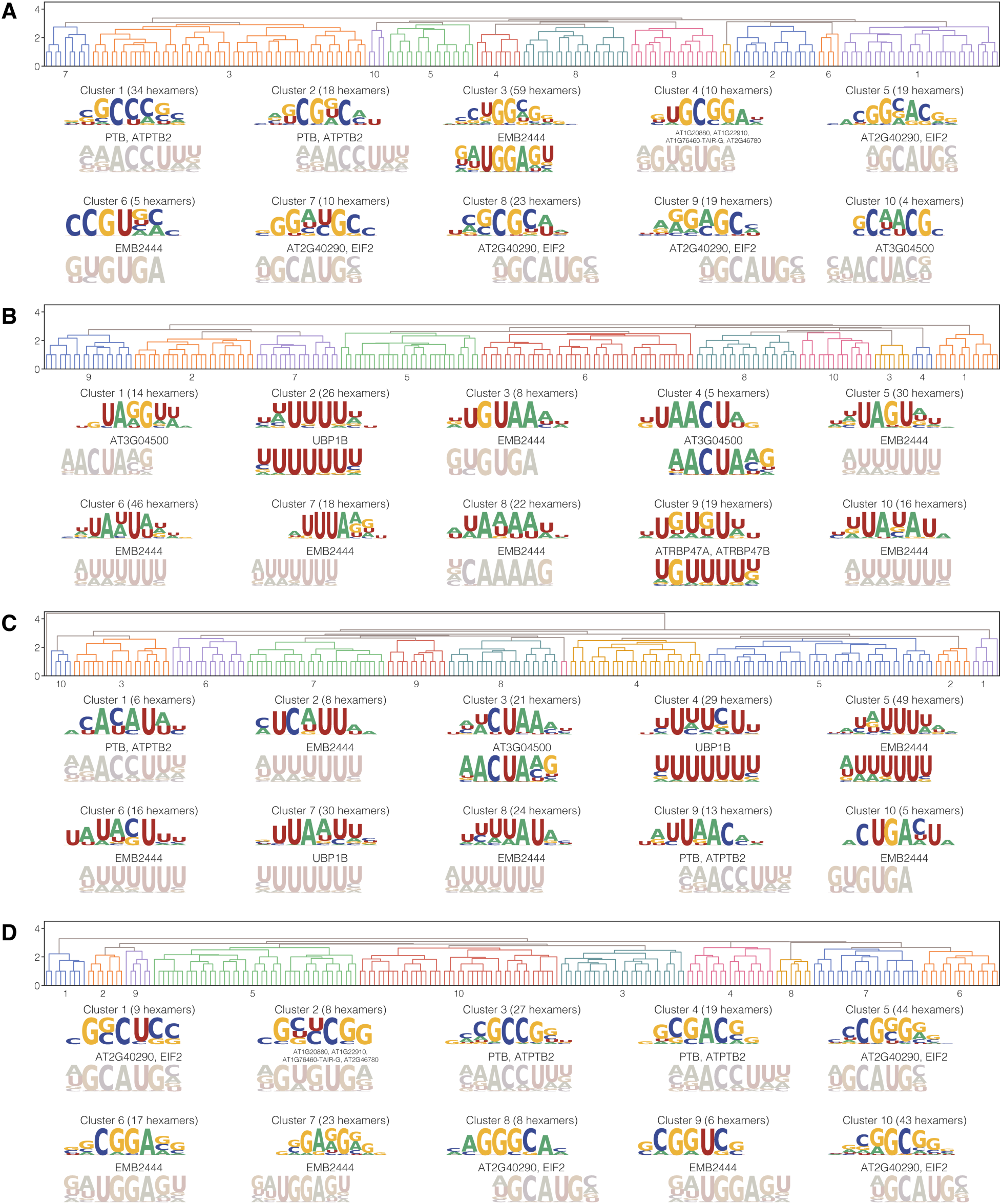
Comparison of *Arabidopsis* SRE motifs to *Arabidopsis* RNACompete motifs. A) Dendrogram for clustering hexamers with the 5% most positive ESR scores in *Arabidopsis*, as well as the resulting aligned clusters and best RNAcompete matches below those clusters. Desaturation indicates that the best match did not satisfy our threshold. B) Dendrogram for clustering hexamers with the 5% most negative ESR scores in *Arabidopsis*, as well as the resulting aligned clusters and best RNAcompete matches below those clusters. C) Dendrogram for clustering hexamers with the 5% most positive ISR scores in *Arabidopsis*, as well as the resulting aligned clusters and best RNAcompete matches below those clusters. D) Dendrogram for clustering hexamers with the 5% most negative ISR scores in *Arabidopsis*, as well as the resulting aligned clusters and best RNAcompete matches below those clusters.

**Supplemental Figure 17:**
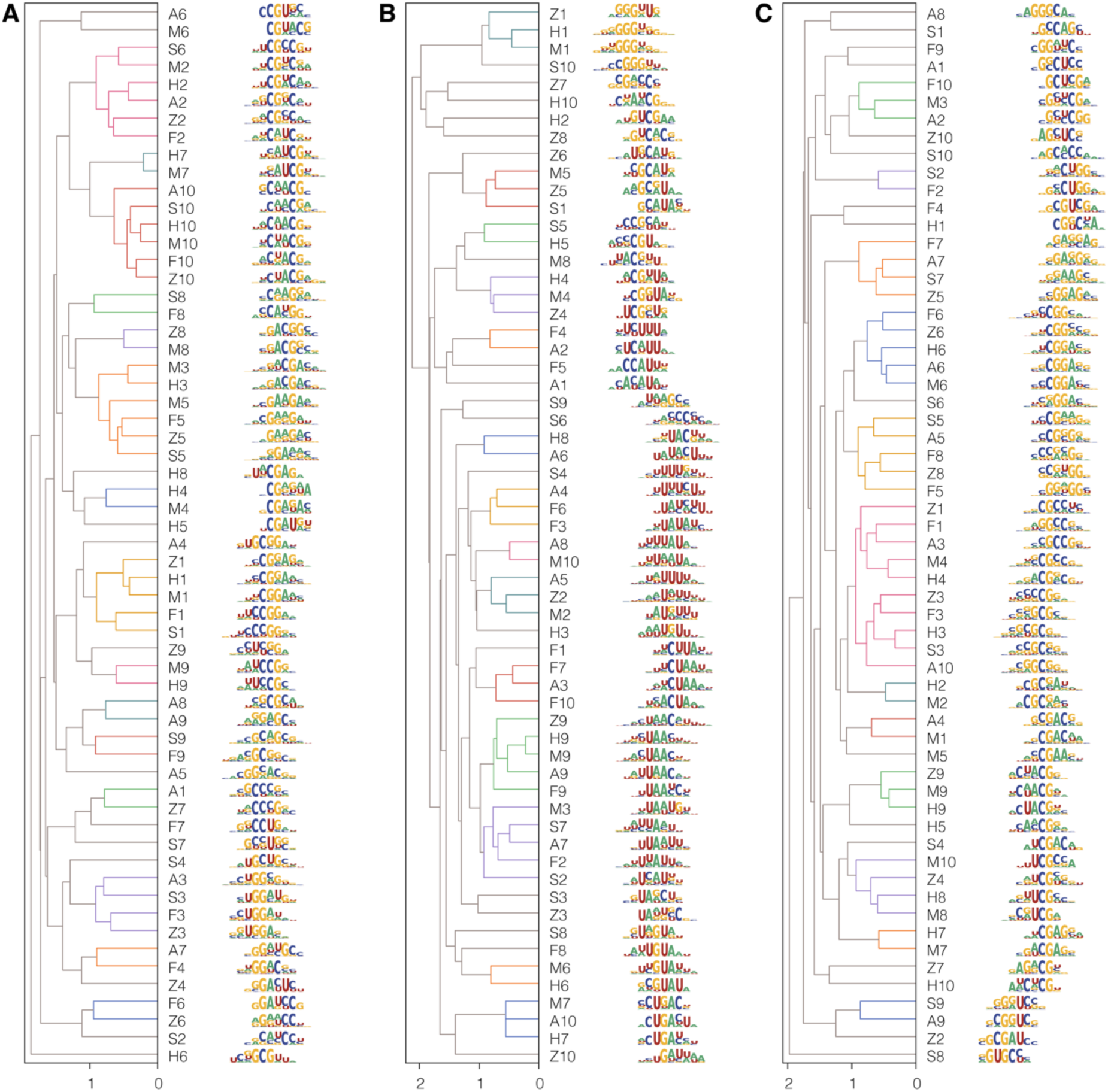
Alignments of ESE, ISE and ISS motifs across organisms. A) All ESE clusters for all organisms clustered by motif similarity. Motifs are labeled using the first letter of the organism, using “S” for silkworm moth, followed by the number of the ESE cluster in Supplemental Figure 7. B) All ISE clusters for all organisms clustered by motif similarity. Motifs are labeled using the first letter of the organism, using “S” for silkworm moth, followed by the number of the ISE cluster in Supplemental Figure 9. C) All ISS clusters for all organisms clustered by motif similarity. Motifs are labeled using the first letter of the organism, using “S” for silkworm moth, followed by the number of the ISS cluster in Supplemental Figure 10.

## Notes

### Competing Interest Statement

The authors have declared no competing interest.

## References

Baeza-Centurion, P., Miñana, B., Schmiedel, J. M., Valcárcel, J., & Lehner, B. (2019). Combinatorial Genetics Reveals a Scaling Law for the Effects of Mutations on Splicing. Cell, 176(3), 549–563.e23. 10.1016/j.cell.2018.12.010

Baeza-Centurion, P., Miñana, B., Valcárcel, J., & Lehner, B. (2020). Mutations primarily alter the inclusion of alternatively spliced exons. ELife, 9, 1–74. 10.7554/eLife.59959

Barash, Y., Calarco, J. A., Gao, W., Pan, Q., Wang, X., Shai, O., Blencowe, B. J., & Frey, B. J. (2010). Deciphering the splicing code. Nature, 465(7294), 53–59. 10.1038/nature09000

Barbeira, A. N., Bonazzola, R., Gamazon, E. R., Liang, Y., Park, Y., Kim-Hellmuth, S., Wang, G., Jiang, Z., Zhou, D., Hormozdiari, F., Liu, B., Rao, A., Hamel, A. R., Pividori, M. D., Aguet, F., Bastarache, L., Jordan, D. M., Verbanck, M., Do, R., … Im, H. K. (2019). GTEx v8 fine mapping on eQTL and sQTL. Zenodo.

Black, D. L. (1995). Finding splice sites within a wilderness of RNA. RNA, 1(8), 763–771.

Busch, A., & Hertel, K. J. (2012). Evolution of SR protein and hnRNP splicing regulatory factors. Wiley Interdiscip Rev RNA, 3(1), 1–12.

Cáceres, E. F., & Hurst, L. D. (2013). The evolution, impact and properties of exonic splice enhancers. Genome Biol, 14(12), R143.

Cheng, J., Nguyen, T. Y. D., Cygan, K. J., Çelik, M. H., Fairbrother, W. G., Avsec, Ž., & Gagneur, J. (2019). MMSplice: modular modeling improves the predictions of genetic variant effects on splicing. Genome Biol, 20(1), 48. 10.1186/s13059-019-1653-z

De Conti, L., Baralle, M., & Buratti, E. (2013). Exon and intron definition in pre-mRNA splicing. Wiley Interdiscip Rev RNA, 4(1), 49–60.

Dominguez, D., Freese, P., Alexis, M. S., Su, A., Hochman, M., Palden, T., Bazile, C., Lambert, N. J., Van Nostrand, E. L., Pratt, G. A., Yeo, G. W., Graveley, B. R., & Burge, C. B. (2018). Sequence, Structure, and Context Preferences of Human RNA Binding Proteins. Mol Cell, 70(5), 854–867.e9. 10.1016/j.molcel.2018.05.001

Fairbrother, W. G., Yeh, R. F., Sharp, P. A., & Burge, C. B. (2002). Predictive identification of exonic splicing enhancers in human genes. Science, 297(5583), 1007–1013.

Fu, X. D., & Ares, M. (2014). Context-dependent control of alternative splicing by RNA-binding proteins. In Nature Reviews Genetics (Vol. 15, Issue 10, pp. 689–701). Nature Publishing Group. 10.1038/nrg3778

Goren, A., Ram, O., Amit, M., Keren, H., Lev-Maor, G., Vig, I., Pupko, T., & Ast, G. (2006). Comparative analysis identifies exonic splicing regulatory sequences–The complex definition of enhancers and silencers. Mol Cell, 22(6), 769–781.

Gupta, K., Yang, C., McCue, K., Bastani, O., Sharp, P. A., Burge, C. B., & Solar-Lezama, A. (2023). Improved modeling of RNA-binding protein motifs in an interpretable neural model of RNA splicing. BioRxiv. 10.1101/2023.08.20.553608

Havens, M. A., & Hastings, M. L. (2016). Splice-switching antisense oligonucleotides as therapeutic drugs. In Nucleic Acids Research (Vol. 44, Issue 14, pp. 6549–6563). Oxford University Press. 10.1093/nar/gkw533

Jaganathan, K., Kyriazopoulou Panagiotopoulou, S., McRae, J. F., Darbandi, S. F., Knowles, D., Li, Y. I., Kosmicki, J. A., Arbelaez, J., Cui, W., Schwartz, G. B., Chow, E. D., Kanterakis, E., Gao, H., Kia, A., Batzoglou, S., Sanders, S. J., & Farh, K. K. (2019). Predicting Splicing from Primary Sequence with Deep Learning. Cell, 176(3), 535–548.e24.

Jens, M., McGurk, M., Bundschuh, R., & Burge, C. B. (2022). RBPamp: Quantitative Modeling of Protein-RNA Interactions in vitro Predicts in vivo Binding. BioRxiv. 10.1101/2022.11.08.515616

Jian, X., Boerwinkle, E., & Liu, X. (2014). In silico prediction of splice-altering single nucleotide variants in the human genome. Nucleic Acids Res, 42(22), 13534–13544. 10.1093/nar/gku1206

Ke, S., Anquetil, V., Zamalloa, J. R., Maity, A., Yang, A., Arias, M. A., Kalachikov, S., Russo, J. J., Ju, J., & Chasin, L. A. (2018). Saturation mutagenesis reveals manifold determinants of exon definition. Genome Research, 28(1), 11–24. 10.1101/gr.219683.116

Keren, H., Lev-Maor, G., & Ast, G. (2010). Alternative splicing and evolution: diversification, exon definition and function. Nat Rev Genet, 11(5), 345–355.

Kumar, S., Suleski, M., Craig, J. M., Kasprowicz, A. E., Sanderford, M., Li, M., Stecher, G., & Hedges, S. B. (2022). TimeTree 5: An Expanded Resource for Species Divergence Times. Mol Biol Evol, 39(8).

Kupfer, D. M., Drabenstot, S. D., Buchanan, K. L., Lai, H., Zhu, H., Dyer, D. W., Roe, B. A., & Murphy, J. W. (2004). Introns and splicing elements of five diverse fungi. Eukaryotic Cell, 3(5), 1088–1100. 10.1128/EC.3.5.1088-1100.2004

Lafferty, J., McCallum, A., & Pereira, F. C. N. (2001). Conditional random fields: Probabilistic models for segmenting and labeling sequence data.

Lim, L. P., & Burge, C. B. (2001). A computational analysis of sequence features involved in recognition of short introns. Proc Natl Acad Sci U S A, 98(20), 11193–11198.

Lorkoví, Z. J., Kirk, D. A. W., Lambermon, M. H. L., & Filipowicz, W. (2000). Pre-mRNA splicing in higher plants. In Reviews (Vol. 5, Issue 4). http://www.cbs.dtu.dk/services/NetGene2/

Mort, M., Sterne-Weiler, T., Li, B., Ball, E. V, Cooper, D. N., Radivojac, P., Sanford, J. R., & Mooney, S. D. (2014). MutPred Splice: machine learning-based prediction of exonic variants that disrupt splicing. Genome Biology, 15(1), R19. 10.1186/gb-2014-15-1-r19

Pai, A. A., Henriques, T., McCue, K., Burkholder, A., Adelman, K., & Burge, C. B. (2017). The kinetics of pre-mRNA splicing in the Drosophila genome and the influence of gene architecture. Elife, 6. 10.7554/eLife.32537

Ray, D., Kazan, H., Cook, K. B., Weirauch, M. T., Najafabadi, H. S., Li, X., Gueroussov, S., Albu, M., Zheng, H., Yang, A., Na, H., Irimia, M., Matzat, L. H., Dale, R. K., Smith, S. A., Yarosh, C. A., Kelly, S. M., Nabet, B., Mecenas, D., … Hughes, T. R. (2013). A compendium of RNA-binding motifs for decoding gene regulation. Nature, 499(7457), 172–177.

Rodriguez, J. M., Maietta, P., Ezkurdia, I., Pietrelli, A., Wesselink, J. J., Lopez, G., Valencia, A., & Tress, M. L. (2013). APPRIS: annotation of principal and alternative splice isoforms. Nucleic Acids Res, 41(Database issue), D110–7.

Rosenberg, A. B., Patwardhan, R. P., Shendure, J., & Seelig, G. (2015). Learning the sequence determinants of alternative splicing from millions of random sequences. Cell, 163(3), 698– 711. 10.1016/j.cell.2015.09.054

Rube, H. T., Rastogi, C., Feng, S., Kribelbauer, J. F., Li, A., Becerra, B., Melo, L. A. N., Do, B. V., Li, X., Adam, H. H., Shah, N. H., Mann, R. S., & Bussemaker, H. J. (2022). Prediction of protein–ligand binding affinity from sequencing data with interpretable machine learning. Nature Biotechnology, 40(10), 1520–1527. 10.1038/s41587-022-01307-0

Sarawagi, S., & Cohen, W. W. (2004). Semi-markov conditional random fields for information extraction. Advances in Neural Information Processing Systems, 17.

Shapiro, M. B., & Senapathy, P. (1987). RNA splice junctions of different classes of eukaryotes: sequence statistics and functional implications in gene expression. Nucleic Acids Res, 15(17), 7155–7174.

Shun-Zheng, Y., & Kobayashi, H. (2003). An efficient forward-backward algorithm for an explicit-duration hidden Markov model. IEEE Signal Processing Letters, 10(1), 11–14.

Sibley, C. R., Blazquez, L., & Ule, J. (2016). Lessons from non-canonical splicing. In Nature Reviews Genetics (Vol. 17, Issue 7, pp. 407–421). Nature Publishing Group. 10.1038/nrg.2016.46

Strauch, Y., Lord, J., Niranjan, M., & Baralle, D. (2022). CI-SpliceAI—improving machine learning predictions of disease causing splicing variants using curated alternative splice sites. Plos One, 17(6), e0269159.

Van Nostrand, E. L., Freese, P., Pratt, G. A., Wang, X., Wei, X., Xiao, R., Blue, S. M., Chen, J. Y., Cody, N. A. L., Dominguez, D., Olson, S., Sundararaman, B., Zhan, L., Bazile, C., Bouvrette, L. P. B., Bergalet, J., Duff, M. O., Garcia, K. E., Gelboin-Burkhart, C., … Yeo, G. W. (2020). A large-scale binding and functional map of human RNA-binding proteins. Nature, 583(7818), 711–719. 10.1038/s41586-020-2077-3

Wahl, M. C., Will, C. L., & Lührmann, R. (2009). The Spliceosome: Design Principles of a Dynamic RNP Machine. In Cell (Vol. 136, Issue 4, pp. 701–718). Elsevier B.V. 10.1016/j.cell.2009.02.009

Wang, Y., Ma, M., Xiao, X., & Wang, Z. (2012). Intronic splicing enhancers, cognate splicing factors and context-dependent regulation rules. Nat Struct Mol Biol, 19(10), 1044–1052.

Wang, Y., Xiao, X., Zhang, J., Choudhury, R., Robertson, A., Li, K., Ma, M., Burge, C. B., & Wang, Z. (2013). A complex network of factors with overlapping affinities represses splicing through intronic elements. Nat Struct Mol Biol, 20(1), 36–45.

Wang, Z., Rolish, M. E., Yeo, G., Tung, V., Mawson, M., & Burge, C. B. (2004). Systematic identification and analysis of exonic splicing silencers. Cell, 119(6), 831–845.

Wieringa, B., Hofer, E., & Weissmann, C. (1984). A minimal intron length but no specific internal sequence is required for splicing the large rabbit beta-globin intron. Cell, 37(3), 915–925.

Wong, M. S., Kinney, J. B., & Krainer, A. R. (2018). Quantitative Activity Profile and Context Dependence of All Human 5’ Splice Sites. Mol Cell, 71(6), 1012–1026.e3. 10.1016/j.molcel.2018.07.033

Xiong, H. Y., Alipanahi, B., Lee, L. J., Bretschneider, H., Merico, D., Yuen, R. K., Hua, Y., Gueroussov, S., Najafabadi, H. S., Hughes, T. R., Morris, Q., Barash, Y., Krainer, A. R., Jojic, N., Scherer, S. W., Blencowe, B. J., & Frey, B. J. (2015). RNA splicing. The human splicing code reveals new insights into the genetic determinants of disease. Science, 347(6218), 1254806. 10.1126/science.1254806

Xiong, H. Y., Barash, Y., & Frey, B. J. (2011). Bayesian prediction of tissue-regulated splicing using RNA sequence and cellular context. Bioinformatics, 27(18), 2554–2562. 10.1093/bioinformatics/btr444

Yeo, G., & Burge, C. B. (2004). Maximum entropy modeling of short sequence motifs with applications to RNA splicing signals. J Comput Biol, 11(2–3), 377–394.

Yeo, G., Hoon, S., Venkatesh, B., & Burge, C. B. (2004). Variation in sequence and organization of splicing regulatory elements in vertebrate genes (Vol. 101, Issue 44). www.pnas.orgcgidoi10.1073pnas.0404901101

